# Identification of long regulatory elements in the genome of *Plasmodium falciparum* and other eukaryotes

**DOI:** 10.1101/2020.06.02.130468

**Authors:** Christophe Menichelli, Vincent Guitard, Rafael M. Martins, Sophie Lèbre, Jose-Juan Lopez-Rubio, Charles-Henri Lecellier, Laurent Bréhélin

## Abstract

Long regulatory elements (LREs), such as CpG islands, polydA:dT tracts or AU-rich elements, are thought to play key roles in gene regulation but, as opposed to conventional binding sites of transcription factors, few methods have been proposed to formally and automatically characterize them. We present here a computational approach named DExTER dedicated to the identification of LREs and apply it to the analysis of the genomes of different eukaryotes including *P. falciparum*. Our analyses show that all tested genomes contain several LREs that are somewhat conserved along evolution, and that gene expression can be predicted with surprising accuracy on the basis of these long regions only. Regulation by LREs exhibits very different behaviours depending on species and conditions. On Apicomplexa organisms, the process appears highly dynamic, with different LREs involved at different phases of their life cycle. For multicellular organisms, the same LREs are involved in all tissues, but a dynamic behavior is observed along embryonic development stages. In *P. falciparum*, whose genome is known to be strongly depleted of transcription factors, LREs appear to be of especially high importance, and our analyses show that they are involved in both transcriptomic and post-transcriptomic regulation mechanisms. Moreover, we demonstrated the biological relevance of one the LREs discovered by DExTER in *P. falciparum* using an *in vivo* reporter assay. The source code (python) of DExTER is available at address https://gite.lirmm.fr/menichelli/DExTER.

## Background

Gene expression is regulated at different levels and by different mechanisms in Eukaryotes. At the DNA level, transcription factors (TFs) are supposed to play a key role by binding specific motifs of typically 6-12bp long in promoters or enhancers. However, TFs are not the only actors, and other mechanisms such as histone occupation, epigenetic marks, transcript stability, 3D structure of the chromatin, etc. are known to be involved in the whole and entangled process of gene expression regulation. Moreover, these mechanisms and their relative importance in the global transcriptomic response are expected to be greatly dependent on species and conditions.

In *Plasmodium falciparum*, the causative agent of malignant malaria in humans, different levels of gene regulation are also present, including cis-regulatory DNA elements, transcriptional factors, epigenetic regulation, and post transcriptional and translation control. Recently, around 4000 regulatory elements have been identified by directly profiling chromatin accessibility. The vast majority of these sites are located within 2 kb upstream of genes and their chromatin accessibility pattern correlates positively with abundance of mRNA transcripts (1). Main factors of the general transcription machinery are present in the plasmodium genome, yet only a few specific plasmodium TFs have been identified and validated. They constitute approximately 1% of all protein-coding genes (2; 3) compared to ∼3% in yeast or 6% in human. Especially, the importance of the apicomplexan AP2 TF family has become apparent in regulating *P. falciparum* biology (4; 5; 6; 7; 8; 9). Among the mechanisms for epigenetic regulation, covalent histone modifications are the best described so far, and experimental evidences show that this form of regulation is most evident in heterochromatin-mediated silencing of genes located in subtelomeric regions and a few chromosome-internal heterochromatic islands while the largest part of the genome is in an euchromatic transcriptionally permissive state (10; 11; 12; 13). Finally, several studies have shown that post-transcriptional regulation (mRNA degradation) and translational control mechanisms also operate in this parasite (see for example (14; 15; 16; 17)).

Study of the links between DNA and gene expression has a long history in bioinformatics. Notably, numerous approaches have been proposed to identify TF motifs by searching for motifs shared by sequences associated with a given gene expression profile (18; 19; 20; 21; 22; 23; 24). In recent years, it has been shown that transcription factor binding but also gene expression as well as several related chromatin features such as histone modifications or DNase I–hypersensitive sites can be predicted from DNA sequence only, often with surprisingly high accuracy (25; 26; 27; 28; 29; 30). With the exception of a few approaches (*e.g.* (26; 30)), deep learning, and particularly convolutional neural networks (CNN), are often used for this task (25; 31; 27; 32; 29). The good predictive performances of these approaches suggest that a large part of the instructions for gene regulation lie at the level of the DNA. However, identifying the exact DNA features captured by CNNs and assessing their respective predictive power remains a difficult task (33). Interesting methods are being developed to post-analyze and interpret learned CNNs in order to identify the DNA determinants used for the predictions (see *e.g.* (25; 34; 33)) but, to the best of our knowledge, these attempts are limited to the identification of single nucleotides and motifs. Besides conventional TFBS motifs, which usually display strong positional information and relatively short length (dozens bp), several studies have highlighted the role that longer regions without clear motif but with biased nucleotide composition can have for regulating gene expression. The most famous example in vertebrates are CpG islands, which are defined as regions larger than 200bp with a high ratio of CpG dinucleotides. CpG islands often correspond to transcription initiation sites (35). While the exact mechanism by which they acquire their function remains a debated subject, they are now widely considered as important regulatory structures of mammalian genomes (36). Similarly, other works have shown that large (hundreds of bp) CpG-rich domains directly downstream of TSS and that do not classify as CpG islands increase transcription rates of endogenous genes in human cells (37). Short tandem repeats are another class of biased regions that could act as regulatory elements. These elements are made of periodic k-mers of 2-6 bp, spanning regions whose total length has been shown to widely impact gene expression and to contribute to expression variation, independently of their genomic location (exon, intron, intergenic) (38). PolydA:dT tracts have been shown to act as promoter elements favoring transcription by depleting repressive nucleosomes (39), specifically by orientating the displacement of nucleosomes (40). Other examples are the AU-rich elements, which are 50-150nt sequences, rich in adenosine and uridine bases. They are located in the 3’-UTRs of many short half-life mRNAs and are believed to regulate mRNA degradation by a mechanism dependent on deadenylation (41). Recently, high-resolution chromatin conformation capture (Hi-C) experiments have revealed the existence of contiguous genomic regions with high contact frequencies referred to as topologically associated domains (TADs) (42). It has been shown that TADs actually correspond to different isochores (*i.e.* large regions with homogeneous G+C content) (43), and that genes within the same TAD tend to be coordinately expressed (44), thus highlighting the role nucleotide composition of large regions may have on the regulation of gene expression. In accordance with these observations, we have shown that gene expression in humans can be predicted with surprising accuracy only on the basis of di-nucleotide frequencies computed in predefined gene regions (close promoters, upstream and downstream promoter regions, 5’- and 3’-UTRs, exons, introns) (26). Importantly, we observed that although CpG content in promoters has high contribution when predicting gene expression, dinucleotides other than CpG are also important and likely contribute to gene regulation (26). In line with these works, Quante and Birds have previously proposed that domains with specific base compositions might modulate the epigenome through cell-type-specific proteins that recognize frequent, short k-mers (45). Likewise, Lemaire et al. have found that exon nucleotide composition bias establishes a direct link between genome organization and local regulatory processes, like alternative splicing (46). However, while numerous computational approaches have been developed to identify motifs, no methods have been designed for characterizing these long regulatory regions associated with specific nucleotide composition. Especially, *in silico* methods are needed to automatically identify the boundaries and nucleotide specificities (nucleotide, dinucleotide or longer k-mers) of the long regulatory elements (LREs) present in sequenced genomes. It is worth noting that segmentation methods that aim to identify sections of homogeneous composition along the genome have been proposed, for example for identifying CpG islands (47; 48; 49). However, with these approaches the segmentation is done on the basis of the sequence alone, without using gene expression to help in the segment definition. As a consequence, the inferred regions are not specifically linked to gene expression regulation, and the approach can miss several regulatory elements.

Here we propose a new method named DExTER (Domain Exploration To Explain gene Regulation) for identifying the precise boundaries and nucleotide specificities of long regulatory elements. Given a set of sequences (one for each gene) and gene expression data in a given condition (treatment, time point or cell type), DExTER identifies pairs of (k-mer,region) for which there is a correlation between gene expression and the frequency of the k-mer in the defined region of each gene. DExTER uses an iterative procedure to explore the space of (k-mer,region) by gradually increasing the size of the k-mers. The identified pairs form a set of predictive variables that are then combined to predict gene expression. The predictor is trained with machine learning algorithms based on penalized likelihood that allows us to select a minimal number of predictive variables and hence to identify the most important regions and k-mers that could act as regulatory elements.

We applied DExTER on 4000bp sequences centered around the gene start (most upstream TSS or AUG) of *P. falciparum* and several other organisms in the Eukaryote tree. Depending on the species, the method identified different large regions (hundred of bps) whose enrichment in certain k-mers was correlated with gene expression. We hypothesized that these long biased sequences could constitute regulatory elements, different from the classical TF binding sites that usually involve only a dozen base pairs. For most tested species, we show that these long regulatory elements are predictive of expression with an accuracy in between 50% and 60%. For *P. falciparum* the accuracy even exceeds 70%, indicating that such regulatory elements could have a predominant role in this species. Furthermore, our analysis showed that this regulation is highly dynamic, with different regions and k-mers involved at different stages of *Plasmodium* life cycle. For species outside of the Apicomplexa phylum, the mechanism appears much more static, except in embryonic development of *Drosophila* and *C. elegans*. Further analysis in *Plasmodium* showed a clear dichotomy between identified regulatory elements, with elements located upstream of the TSS being mainly associated with transcriptional regulation, while downstream elements being mainly involved post-transcriptionally. Finally, in order to validate one of the identified elements in *P. falciparum*, a GFP reporter assay was performed, confirming our *in silico* results, thus demonstrating the high importance of this LRE in *P. falciparum* biology.

## Results

### An in silico approach for identifying long sequences linked to expression

The DExTER method takes gene expression data and a set of sequences (one for each gene) aligned on a common anchor. In the experiments below, we took the 4000bp sequences centered either around the gene start (*i.e.* most upstream TSS) or around the AUG but other alignment anchors could also be used (gene end, exon-intron boundaries, etc.). In the first step (feature extraction), DExTER identifies pairs of (k-mer,region) for which the frequency of the k-mer in the defined region is correlated with gene expression. Sequences are first segmented in different bins. We used 13 bins in the following experiments. The number of bins impact the precision of the approach but also the computing time of the analysis. DExTER starts with 2-mer (dinucleotides) and, for each 2-mer, identifies the region of consecutive bins for which the 2-mer frequency in the region is mostly correlated with gene expression. A lattice structure is used for this exploration (see Figure 1 and details in section Methods). Once the best region has been identified for a 2-mer, DExTER attempts to iteratively extend this 2-mer for identifying longer k-mers. For each considered k-mer, a lattice analysis is run to identify the best region associated with this k-mer. At the end of the process, a set of variables corresponding to the frequency of the identified k-mers in the identified regions are returned for each gene. In the second step, DExTER learns a model that predicts gene expression on the basis of these variables. We used a linear regression model:

**Figure 1.**
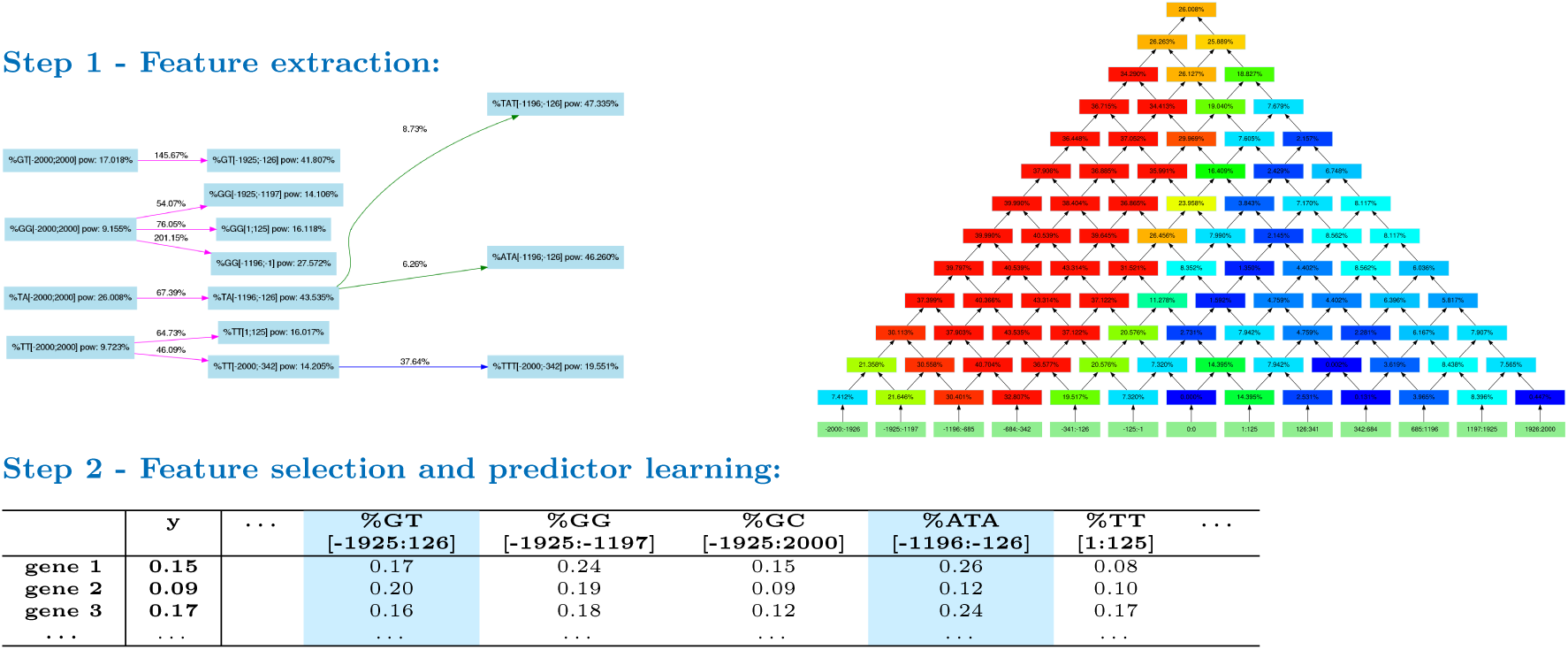
The DExTER method. In step 1, DExTER attempts to identify pairs of (k-mer,region) for which the frequency of the k-mer in the defined region is correlated with gene expression. DExTER starts with a 2-mer and compute a lattice (right) representing different regions. The top of the lattice represents the whole sequence, while lower nodes represent smaller regions. At each position, the correlation between 2-mer frequency and gene expression is computed, and regions with highest correlation are identified. Then, the 2-mer is extended to 3-mers, and the correlation with expression are computed in the best regions. If the correlation increases, the whole process is repeated with increasing k-mers. Otherwise, DExTER starts a new exploration from a different 2-mer, until every 2-mer has been explored. This way, different variables (*i.e.* pairs of (k-mers-regions)) are iteratively built (see the exploration graph on the left). In step 2, the frequency of all variables identified in step 1 are gathered into one long table. Then, a linear model predicting gene expression from a linear combination of the variables is learned. A special penalty function (LASSO) is used during training, for selecting only the best variables in the model (blue columns). If several gene expression data are available for one species (*i.e.* several *y* vectors), then step 1 is ran independently on each data, and all identified variables are gathered into a single table. Then, a linear model is learned for each data, but the different models are learned simultaneously with another penalty function that tends to select the same variables for the different data (group LASSO for multitask learning, see Methods).

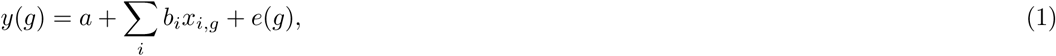

where *y*(*g*) is the expression of gene *g, x*_*i,g*_ is variable *i* for gene *g, e*(*g*) is the residual error associated with gene *g, a* is the intercept and *b*_*i*_ is the regression coefficient associated with variable *i*. Because the set of variables identified in the first step may be large and variables are often correlated, the model is trained with a lasso penalty function (50) that selects the most relevant variables solely (feature selection). Finally, once a model has been trained, its accuracy is evaluated by computing the correlation between predicted and observed expressions on several hundred genes. To avoid any bias, this is done on a set of genes that have not been used in the two previous steps.

### Long sequences with specific composition are predictive of expression for several Eukaryotes and especially for *P. falciparum*

The approach has been applied to several series of expression data, targeting unicellular and multicellular eukaryotes in different conditions. Besides the erythrocytic cycle of *P. falciparum* (51), we also studied the *P. berghei* life cycle (52) as well as that of *T. gondii* (53), another species in the Apicomplexan taxa to which belong the two *Plasmodium*s. The *S. cerevisiae* cell cycle (54) completes the comparisons for unicellular organisms. For multicellular organisms, two series monitoring tissues and development were analyzed for *Drosophila* (55; 56), *C. elegans* (57; 58), human (59; 60), and the plant *A. thaliana* (61; 62). Note that one model is learned for each condition, hence several models are learned for each series using multitask learning (see methods). In all experiments, 2/3 genes are used for training (Step 1 and 2) and 1/3 genes are used for measuring accuracy (the same training/testing gene sets are used in all conditions of the same series). Bar charts on Figure 2 summarizes the accuracy achieved in the different conditions. Except for *P. falciparum* and *P. berghei*, all sequences are centered around the gene start (*i.e.* most upstream TSS). For *Plasmodium* species, we obtained higher accuracies with sequences centered on the AUG, so we used this anchor in the experiments below.

**Figure 2.**
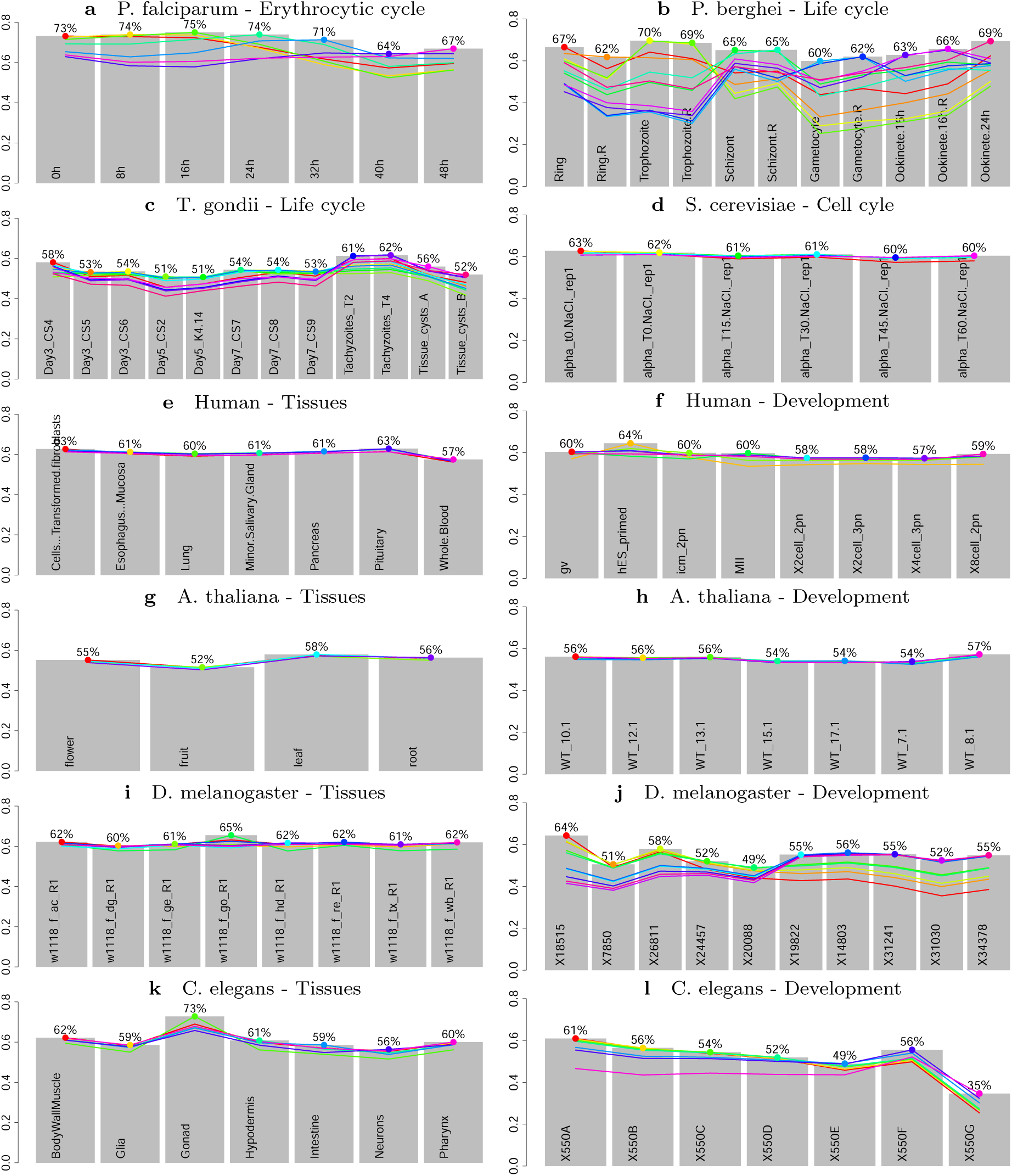
Accuracy of the DExTER models for predicting coding-gene expression in different species and conditions. Grey charts represent the accuracy, measured as the correlation between predicted and observed gene expression, of the models learned on different conditions. Colored curves summarize the accuracy of a model learned on a specific condition (identified by a big dot of the same color) when used to predict the other conditions of the same series.

The accuracy of the method fluctuates around 60% for most species, thus generalizing our previous study on human that showed that gene expression can be predicted with surprisingly high accuracy using only nucleotide frequencies of specific large gene regions (26). Intriguingly, for *P. falciparum* the accuracy exceeds 70% on many stages, which is of particular resonance in an organism for which most attempts for identifying TFs have been unsuccessful.

We first asked whether these long regions detected by our approach may correspond to multiple occurrences of classical TF binding motifs such as the ones referenced in databases like TRANSFAC (63) and JASPAR (64). To do so, we concentrated on the 5 most important variables identified by the learning procedure in each condition (see Methods for details). Figure 3 reports the distribution of the lengths of k-mers and regions of these variables, as well as the distribution of the median number of k-mer occurrences in the identified region of the sequence. In most cases, the large size of the regions (hundreds base pairs), the shortness of the k-mers (3 or less) and the high number of occurrences (median number > 20) seem incompatible with classical TFBSs, which usually involve a dozen base pairs and, to the best of our knowledge, are not known to repeat such high number of times on such long regions. Actually, from the 154 studied variables, we estimate that less than a dozen may correspond to traditional TFBS motifs. A notable exception is the k-mer AGACA identified in *P. berghei*, and whose frequency in the identified region is either 1 or 0 for most sequences. Apart from other variants of this k-mer (which represent most of the identified exceptions) one can also distinguish the short motif TTA, whose presence at the exact position of gene TSSs in *C. elegans* seems negatively correlated with expression.

**Figure 3.**
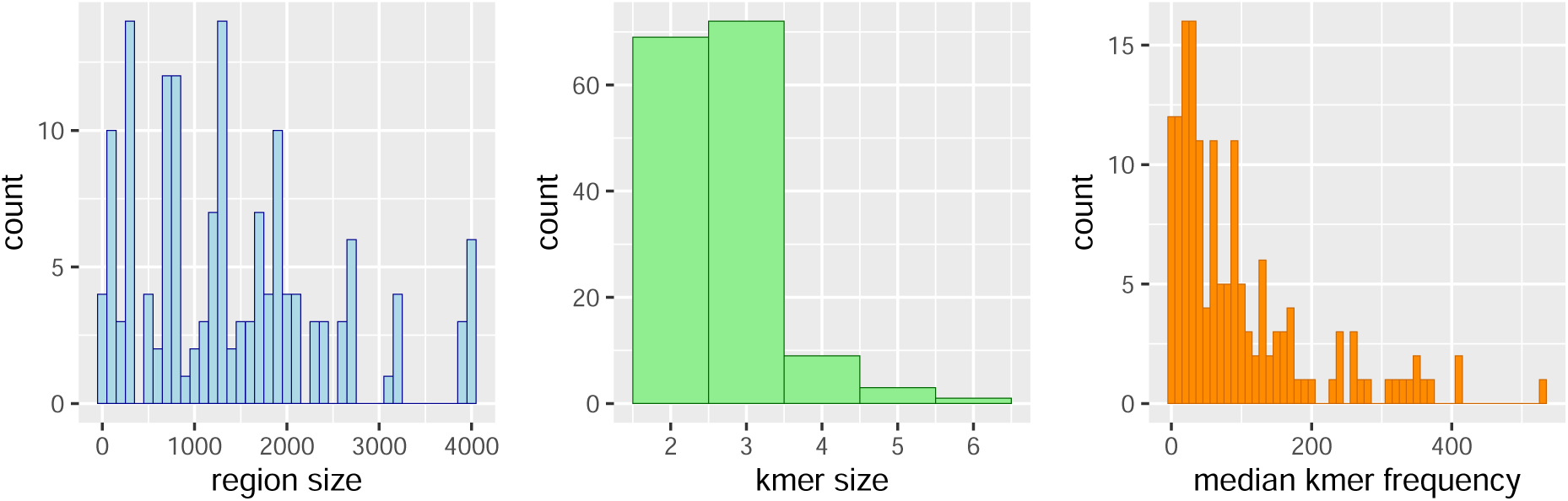
Lengths and frequencies of the variables identified in the different species and conditions. The left histogram reports the distribution of k-mer lengths of the most important variables identified in all species and conditions, while the middle histogram reports the distribution of region lengths of these variables. The right histogram reports the median number of occurrences of the identified k-mers in the identified regions in all studied sequences of the different species.

We then studied more closely the way k-mer occurrences are distributed along the identified regions. We checked whether the occurrences tend to appear isolated or inside repeat blocks. We analyzed for this the most important variable of each species. Supp. Figure 1 reports for each species and each gene the proportion of occurrences that appear isolated, while Supp. Figure 2 reports the histogram of repeat block length. For these analyses, a repeat block involves at least two k-mer occurrences that are either immediately consecutive or overlapping (for example, sequences ATAATA and ATATA are two blocks made up of two repeats of the ATA k-mer). Except for the two represented *Plasmodium* variables, more than 75% of k-mer occurrences are isolated, and the few repeat blocks are mainly made up of two repeats. The picture is different for the two variables in *Plasmodium* species, as isolated occurrences seem much rarer (around 25%), and the size of a typical repeat block usually fluctuates in between 2 and 20 repeats.

### Dynamics, composition, and location of long regulatory elements differ depending on species and conditions

We next sought to test whether LREs are associated with dynamic or static regulation processes. For this, each model learned in a specific condition was used to predict expression in other conditions of the same series, and accuracy was measured. Colored curves on Figure 2 summarize these permutation experiments. Static and dynamic behaviors appear to coexist and to be highly dependent on species and conditions. While approximately the same model is learned on the different tissues of human, *A. thaliana, Drosophila* and *C. elegans*, in the two *Plasmodium*s a model learned on a specific stage has poor accuracy on the other stages, even when the permutations are restricted to the erythrocytic cycle. For *T. gondii* the behavior is similar, although much less strong, while for *S. cerevisiae* the mechanism seems completely static. Interestingly, a dynamic behavior is also observed on developmental series of *C. elegans* and *Drosophila* although almost no differences can be observed when permuting the models learned on different tissues of these organisms. Note however that, for both species, slight differences can be observed in gonads, and that this tissue is also the one were the accuracy is the highest. Part of differences between species can be explained by the inherent correlation of expression between conditions, which are, on average, higher in Human, *Drosophila, C. elegans, A. thaliana* and yeast than in *Plasmodium*s and *T. gondii* (see Supp. Figure 3). However, this does not seem to be the only reason for the static behavior observed in former species. Indeed, when we restrict the comparisons to pairs of conditions with similar expression correlations, *Plasmodium* models are still more different than tissue models of Human, *Drosophila* or *C. elegans*. For example, the 0h/48h *P. falciparum* pair, and the whole-blood/pituitary human pair have both expression correlation around 80% but, contrary to *P. falciparum*, the human models seem completely interchangeable. Similarly, several pairs of tissues of *A. thaliana* and *C. elegans* show only moderate expression correlations, although approximately the same model is learned on these tissues.

We next sought to compare the composition and location of LREs identified in the different species and conditions. For this, we concentrated on the 5 most important variables identified by the learning procedure in each condition. Figure 4 reports the correlations between these variables and expression in the different conditions. In accordance with the above permutation experiments, we can observe that for *P. falciparum, P. berghei, T. gondii*, as well as for *Drosophila* and *C. elegans* development series, the correlation between variables and expression fluctuates along with the conditions, while for the other series correlations are steadier. For *P. falciparum* we can even observe a sinusoidal behavior, highly reminiscent of the sinusoidal pattern of expression observed in the different studies monitoring gene expression during the erythrocytic cycle (65; 66; 67). We can also observe that the locations of LREs are diverse, depending on species and conditions. In the following, we assigned the variables to six different genomic regions: distal and close promoter, center region, 5’UTR, gene body, and whole region (see Methods for details about how variables were associated with one of these locations). Then, for each variable, we computed a usage statistics that summarizes its importance in the learned models (see Methods). Figure 5 reports the relative importance of the 6 regions in the different conditions (see also Supp. Figure 4 which summarizes the variable usage in the different conditions). Some interesting trends emerge from this analysis. For example, we can observe the importance of distal promoter regions in *P. falciparum* compared to all other species. In human, *Drosophila* and *C. elegans*, center and/or 5’UTR variables are important, especially for tissues. Interestingly, these variables seem less important in gonads and at early time points of *Drosophila* and *C. elegans* development, but take increasing importance along the course of the development of these species. Similarly, for *P. falciparum* the role of promoter variables decreases along the phases of the erythrocytic cycle.

**Figure 4.**
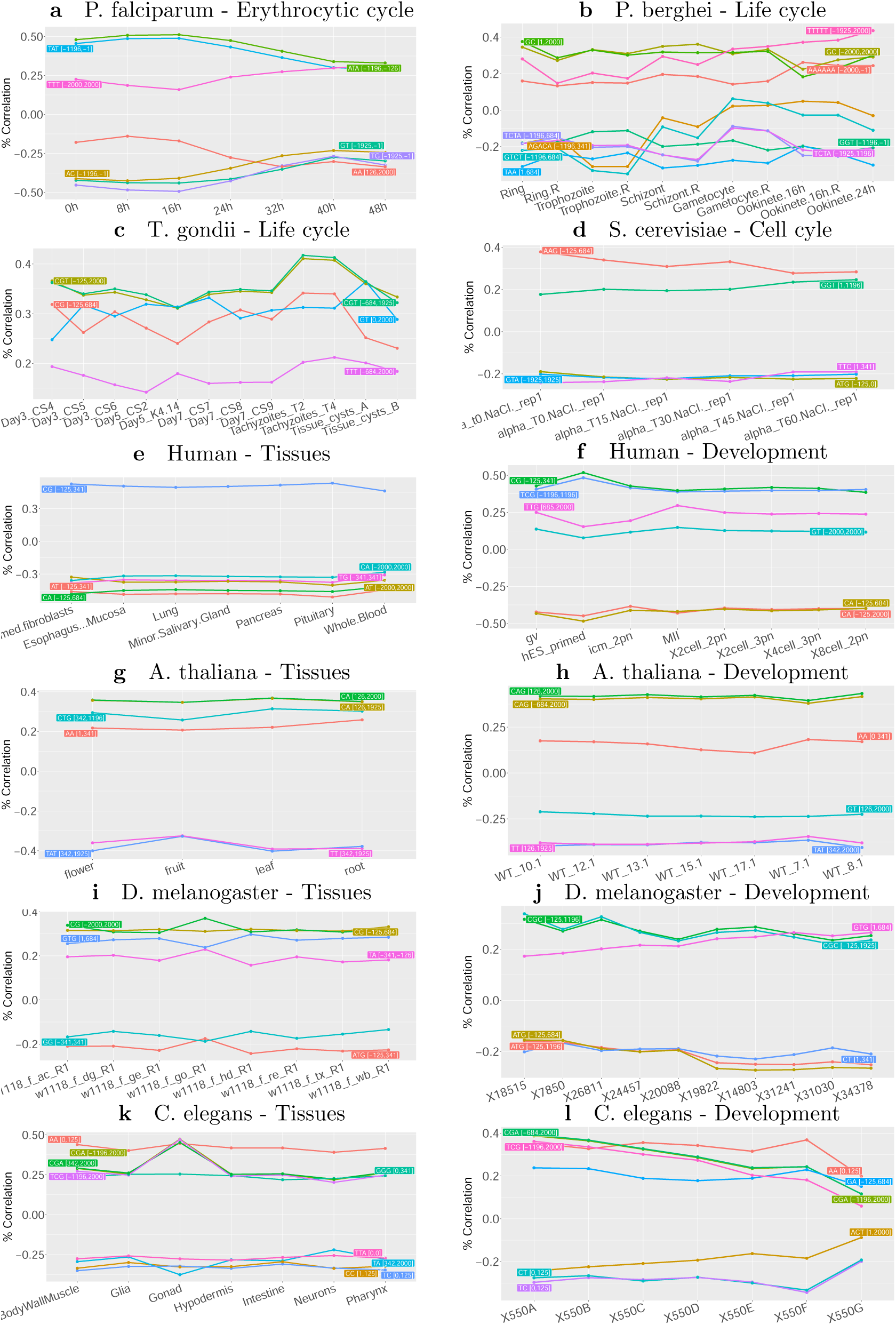
Correlations between expression and k-mer frequency of the most important variables identified in the different species and conditions. For each expression series, the 5 most important variables of each condition were identified, and their correlation to expression were computed for all conditions of the series. Note that there are often more than 5 variables in these figures because the 5 most important variables may vary depending on conditions.

**Figure 5.**
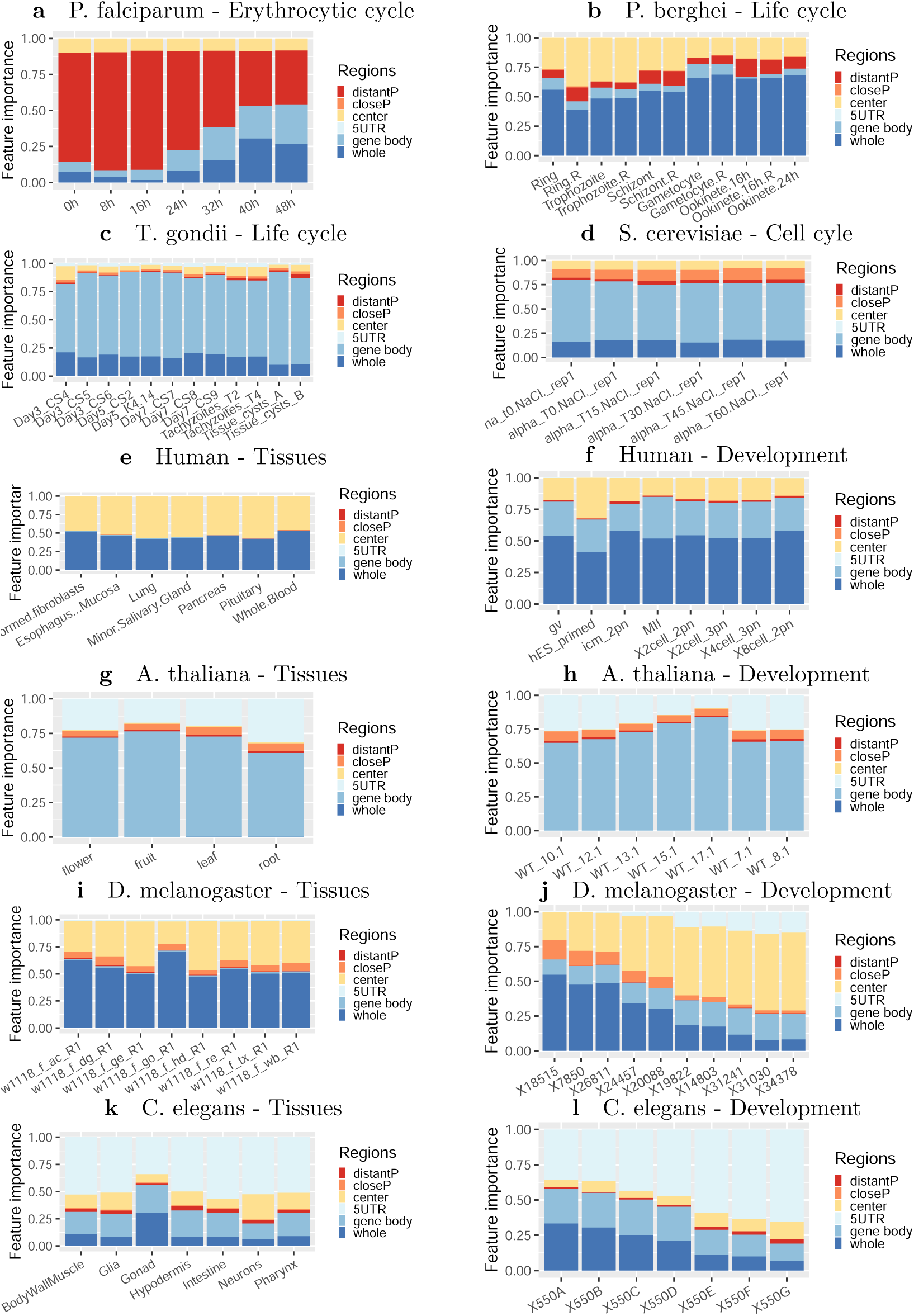
Relative importance of promoter, untranslated and coding regions for predicting gene expression in different species and conditions. For each condition, the 30 most important variables of the model were identified and a usage statistic reflecting the importance of the variables for the prediction was computed (see Method). Then, each variable was associated with one gene region (6 different regions were considered: distal and proximal promoters, center, 5’UTR, gene body, or whole; see Method), and the usage statistics of the variables that belong to the same region were cumulated.

Finally, as a first attempt to assess the conservation of the identified LREs along evolution, we gathered the most important variables identified in each species and conditions, and computed the correlations between these variables and expression in every species and conditions. Then, an unsupervised clustering was run to classify the conditions according to these correlations (see Figure 6a). As we can see, conditions can be perfectly classified on the basis of these correlations (all conditions related to the same species group together). Interestingly, *P. falciparum* and *P. berghei* conditions also group quite clearly together and, with a less clear signal, with *T. gondii* conditions, while the remaining of the groupings does not seem to be in accordance with the phylogenetic tree of Eukaryotes. Looking at the correlation conservation of each variable individually gives a more precise view of this general trend (Figure 6b). Several variables are correlated with expression for both *P. falciparum* and *P. berghei* (*e.g.* ATA [-1196,-126]). At the *Apicomplexa* level, the number of common variables is low but still present (*e.g.* TTT [-684,2000]). Similarly, some variables are common to *Drosophila* and *C. elegans*, and a few ones seem common to *Drosophila, C. elegans* and human (CG [-125,1196]).

**Figure 6.**
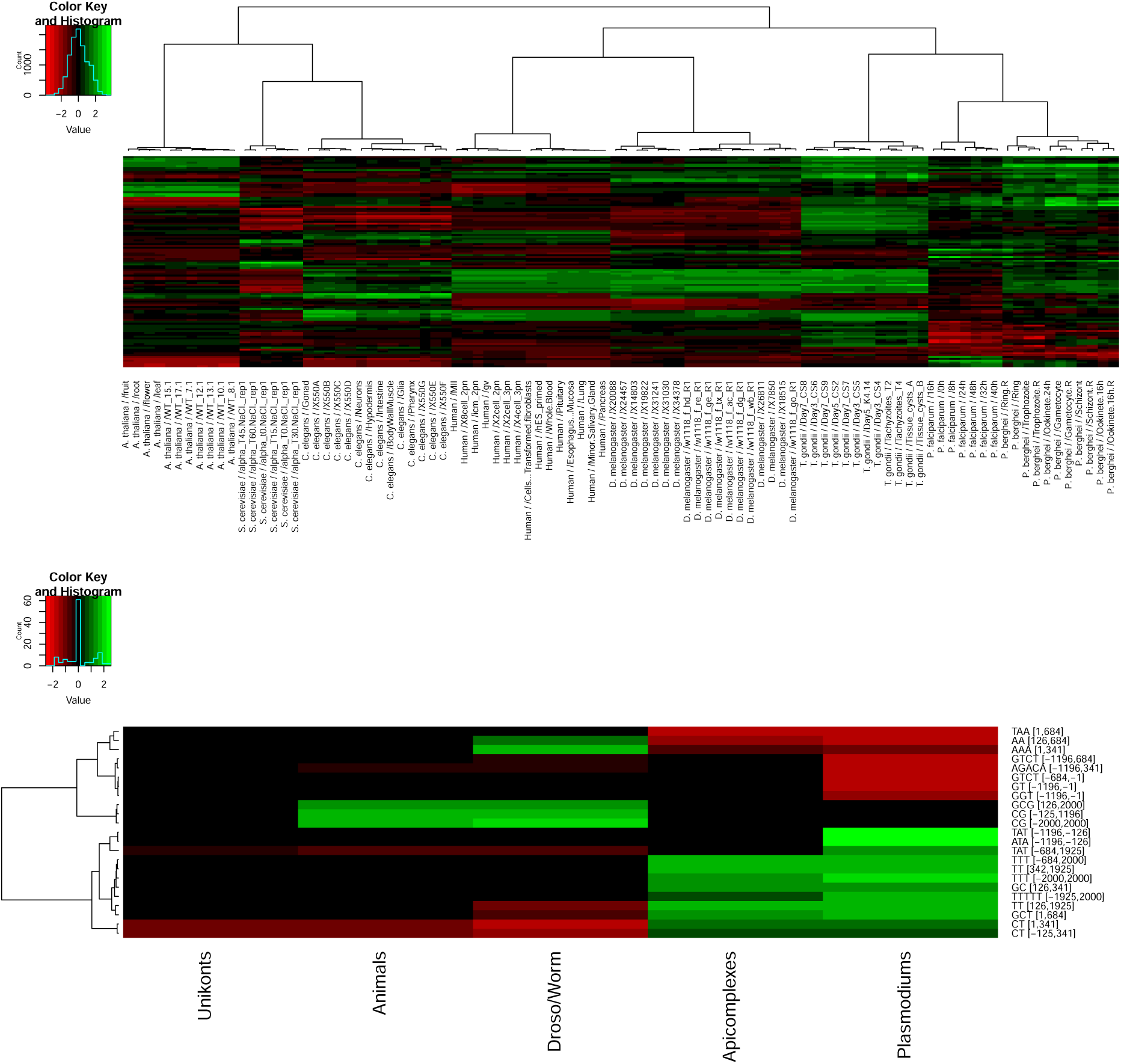
Conservation of long regulatory elements along evolution. The 10 most important variables of each species and conditions were identified and collected, and their correlations with expression were computed for every species and conditions. Correlations were then normalized by conditions (i.e. correlations were divided by the standard deviation of all correlations computed for the condition) to get the same range of values for each condition. **(up)** A hierarchical clustering (Ward’s criterion) was run to classify the conditions according to these correlations. **(down)** The heatmap represents the variables whose correlation with expression is conserved at the level of at least one of five different taxa. The variables that do not show conservation of correlation at any of the 5 taxa have been removed for readability.

### Long regulatory elements are associated with highly dynamic regulations along *P. falciparum* life cycle and have specific GO terms

As explained above, the accuracy of the predictions is especially high for *P. falciparum*, culminating to 74% in the first stages of the erythrocytic cycle. For comparison, we trained several deep learning models (CNNs) on the same *P. falciparum* data. We used for this an architecture similar to that used in DeepSea (25) (see Methods and Supp. Figure 5). The accuracy achieved in these experiments is lower than that obtained with DExTER, but the same dynamic behavior is observed, indicating that CNNs can also capture, at least partially, LRE effects (see section Discussion for possible explanations of the differences observed between CNNs and DExTER). Although LREs can be identified in all studied Eukaryotes, the higher accuracy in *P. falciparum*, along with its dynamic behavior suggests that LREs are particularly important for gene expression regulation in this species. Hence, *P. falciparum* appears as a model of choice for studying the regulatory mechanisms associated with such sequences.

To measure the extent to which LREs control gene expression along the whole life cycle of *P. falciparum*, we ran an analysis of the data of Lopez-Barragan et al. (68) that measures gene expression in sexual and asexual stages of the parasite. Results are summarized in Figure 7. They are globally concordant with those achieved on data exclusively targeting the erythrocytic cycle, with accuracy above 70% in several stages. What is striking, however, is the highly dynamic behavior of the regulation process, something already observed on *P. berghei* life cycle (see Figure 2b): a model with high accuracy on gametocytes has very poor accuracy in asexual stages (particularly in ring stage), and reciprocally. This can be also observed by the high fluctuations of correlation between variable frequencies and gene expression (Figure 7c). Among the best variables identified by DExTER at the different stages, several ones are similar to those identified in the data of Otto et al. (51) (for example ATA and TG on upstream sequences, or the T repeat on whole sequences). Some others seem more correlated with expression in sexual stages than in asexual stages. For example TATAT in [-1196,1925] fluctuates between 30% and 50% correlation, while ATTA[-1925,1925] fluctuates between 0% and −40% correlation. On the whole, upstream variables seem highly important at the beginning of asexual stages, but much less in gametocytes and ookinetes (Figure 7b).

**Figure 7.**
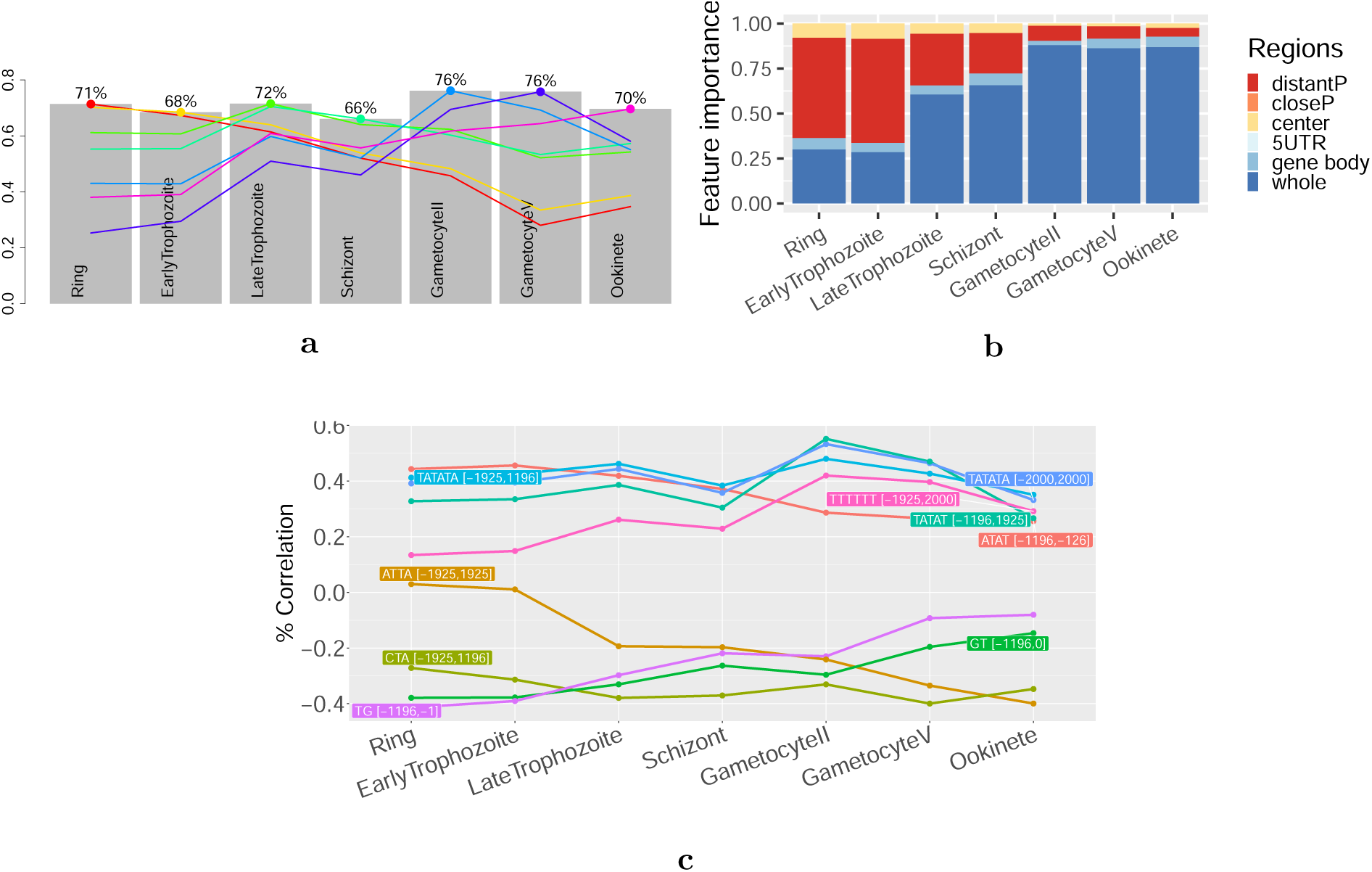
Importance of LREs along the whole life of P. falciparum. **a** Grey charts represent the accuracy, measured as the correlation between predicted and observed gene expression, of the models learned on different phases of *P. falciparum* life cycle. Colored curves summarize the accuracy of a model learned on a specific phase when used to predict gene expression of other phases. **b** Estimate of the importance of upstream, downstream, center and whole regions for predicting gene expression in the different phases. **c** Correlations between expression and k-mer frequency of the 5 most important variables identified at each phase. Because the most important variables vary depending on conditions, the total number of variables is > 5 in this figure.

We next used the GSEA method (69) to analyze some of the variables that show the highest correlation with expression in different phases. Interestingly, genes enriched for specific variables are also associated with specific GO terms (see Supp. Figure 6). For example, genes with high ATA frequency on upstream sequences are associated with high expression in early phases of the erythrocytic cycle and are involved in translation. Genes with high TTT frequency on the whole sequence are highly expressed on later time points and are involved in transport regulation. Similarly, genes with low AA on downstream regions are associated with high expression on late stages and are involved in different metabolic processes. Finally, genes with high TATATA frequency on the whole sequence are more expressed in gametocytes and are involved in chromatin assembly.

### Long regulatory elements are associated with transcriptional and post-transcriptional regulations in the *P. falciparum* intraerythrocytic cyle

We next analyzed more closely the timing of the LREs identified in the intraerythrocytic cycle (IEC). Figure 8a shows the 8 most important variables identified in each stage of the IEC (representing a total of 10 different variables). Left and right heatmaps present the variables with higher correlation in early (0h-16h) and late (24h-48h) stages of the IEC, respectively. In accordance with results presented in Figure 5, upstream variables are more correlated to expression in early time points, while downstream and whole variables show more correlation in late time points. Next, we estimated the strand specificity associated with each variable. For this, we computed the frequency of the corresponding k-mer in the identified region on the plus strand (as it is done in step 1 of DExTER) and on the minus strand, and we compared the correlations between these two frequencies and expression. Variables for which correlations differ between strands are considered as strand-specific (see Methods for details). In Figure 8a, strand specificity is represented with a color code that goes from blue (no strand specificity) to orange (high strand specificity). Interestingly, all upstream variables show little or no strand specificity, while three among the four variables with highest correlation in late stages are strand-specific.

**Figure 8.**
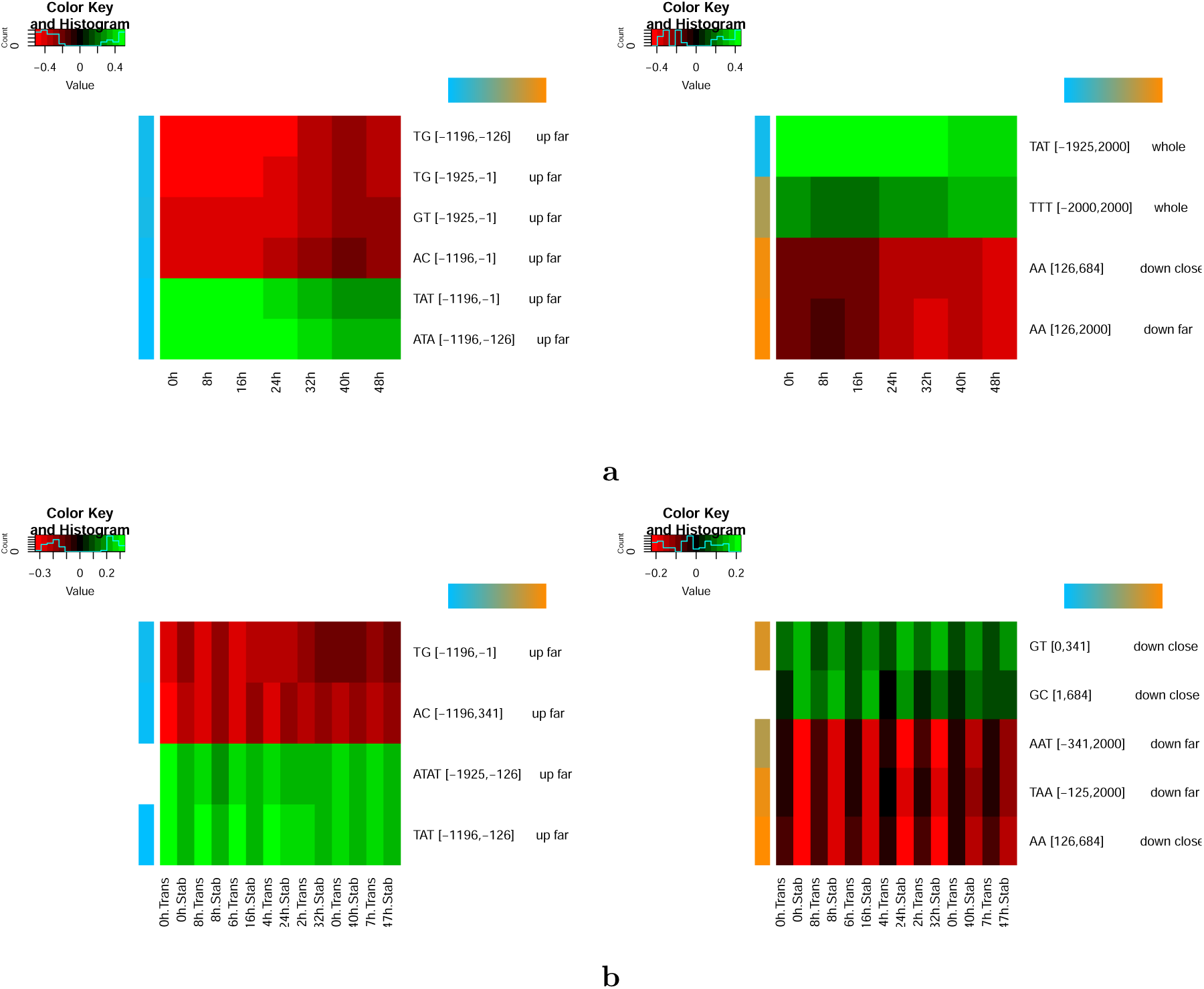
Features identified in the intraerythrocytic cycle of P. falciparum. **a** Heatmaps of correlations between gene expression and most important features identified at each time point of Otto et al. (2010) data. The left heatmap corresponds to features with higher correlation in early time points (0h - 16h), while the right heatmap corresponds to features with higher correlation with late time points (24h - 48h). **b** Heatmaps of correlations between gene expression and most important features identified at each time point of Painter et al. (2018) data. The left heatmap corresponds to features with higher correlation with transcription data, while the right heatmap corresponds to features with higher correlation with stabilization data.

The absence of strand specificity in upstream LREs and its presence in downstream/whole LREs suggest that upstream and downstream LREs may be involved in transcriptional and post-transcriptional regulation mechanisms, respectively. To assess this point, we analyzed the data of Painter et al. (2018) (15), where the level of nascent transcription and stabilized mRNA along the IEC are measured in parallel. We ran DExTER on these two types of data and for each available time point, and identified the 8 most important variables on each condition. Among the 15 different variables, 4 are clearly more correlated with nascent transcription than with stabilized transcript levels, while 5 others are more associated with stabilized transcripts than with nascent transcription (Figure 8b) (the remaining variables cannot be associated clearly with one or the other type of data, see Methods). Remarkably, all variables associated with nascent transcription are both upstream and non strand-specific, while variables associated with mRNA stabilization are downstream and strand-specific.

This raises a question about the nature of the few variables that span the whole sequence. This is typically the case of the variable TTT[-2000,+2000] on IEC (see Figure 4a) and of variable ATTA[-1925,1925] on the whole parasite life cycle (see Figure 7c). We thus measured separately the correlation with expression and the strand specificity of these variables upstream and downstream of the ATG, using time point 48h and the ookinete stage as references. First, for both variables, the correlation with expression is higher for the whole sequence (34% and −37% for TTT and ATTA respectively) than for the upstream (19% and −26%) or downstream (26% and −30%) sequences solely. Second, the strand specificity of these variables is low for upstream sequences (the correlation with expression is almost the same when the variables are computed on the plus or on the minus strand) but high for downstream sequences: the correlation of ATTA with gene expression drops to 8% only, and the correlation of TTT with expression is inverted (−27% on the minus strand *vs.* +27% on the plus strand). Hence, a possible hypothesis for these whole-sequence variables would be that they actually involve two LREs acting coordinately: one upstream LRE associated with transcriptional regulation mechanism, and one downstream LRE likely associated with post-transcriptional regulation. Because the two LREs involve the same k-mer and act in a coordinate way (they are both either positively or negatively correlated with expression) they appear as a single variable in the DExTER analysis.

### Links with histone modifications and variants in *P. falciparum*

Read et al. (70) have shown that gene expression in *P. falciparum* can be predicted with rather good accuracy from various epigenetic marks. Notably, histone variant H2A.Z, and histone modification H3K9Ac and H3K4me3 in promoters and gene bodies appear to be among the most predictive marks for expression. Hence, we sought to assess whether some of the predictive variables identified by our approach could actually be related to these specific marks. To do so, we used the data of Bartfai et al. (71) to compute the H2A.Z, H3K9Ac, and H3K4me3 signals upstream and downstream the AUG codon of every gene, and we ran DExTER to predict these data instead of gene expression. Globally, prediction accuracies are lower than for gene expression. Only H2A.Z and H3K9ac downstream signals can be predicted with accuracy around 60%, but without reaching the > 70% accuracy achieved for gene expression (see Supp. Figure 9). Analysis of the most important variables of the H2A.Z and H3K9ac downstream models shows that several variables identified for gene expression are also singled out when predicting these histone marks (Supp. Figure 10), but none of these variables seems more correlated with histone marks than with gene expression.

### Reporter assay validates a LRE controlling gene expression in *P. falciparum*

Finally, to validate our approach and demonstrate the importance of LREs in *P. falciparum*, a GFP reporter assay was performed. One of the most important variables identified by DExTER is the ATA frequency in region [-1196,-126]. This variable by itself links gene expression and k-mer frequency with nearly 50% correlation at ring stage parasites (Figure 4a). We chose for our analysis the PF3D7 0913900 gene because although the frequency of ATA in region [-1196,-126] is low, it has high ATA content in a very short region [-483,-128]. As expected, this gene is not highly expressed in ring stages (68; 51). We built a chimeric promoter containing 3 repetitions of the region with high ATA frequency [-483,-128] and we named it *chimeric high ATA* (see Figure 9a). As a control, we constructed a second promoter replacing two of the repetitions by a low ATA area that was obtained from the upstream region of the PF3D7 0805300 gene (*chimeric low ATA*, see Figure 9a). Then we used the DExTER model learned at 8h of the erythrocytic cycle (51) to predict the transcriptional activity of both promoters when associated with a GFP gene. DExTER predicted a higher activity for the chimeric high ATA promoter than for the chimeric low ATA promoter. To validate these predictions, each chimeric promoter driving the expression of a reporter GFP gene was integrated in the genome of *P. falciparum* using the CRISPR/Cas9 technology (Ghorbal et al., 2014; see Materials) in the *pfs47* locus (Knuepfer et al., 2017). The transcriptional activity of the chimeric promoters was measured by qPCR analysis of RNA collected at ring stages for each transgenic parasite line. Chimeric high ATA promoter presented a much higher transcriptional activity compared to chimeric low ATA promoter, around 10-folds higher (Figure 9b). These results validate DExTER for identifying LREs, and demonstrate the importance of these elements for the control of gene expression in *P. falciparum*.

**Figure 9.**
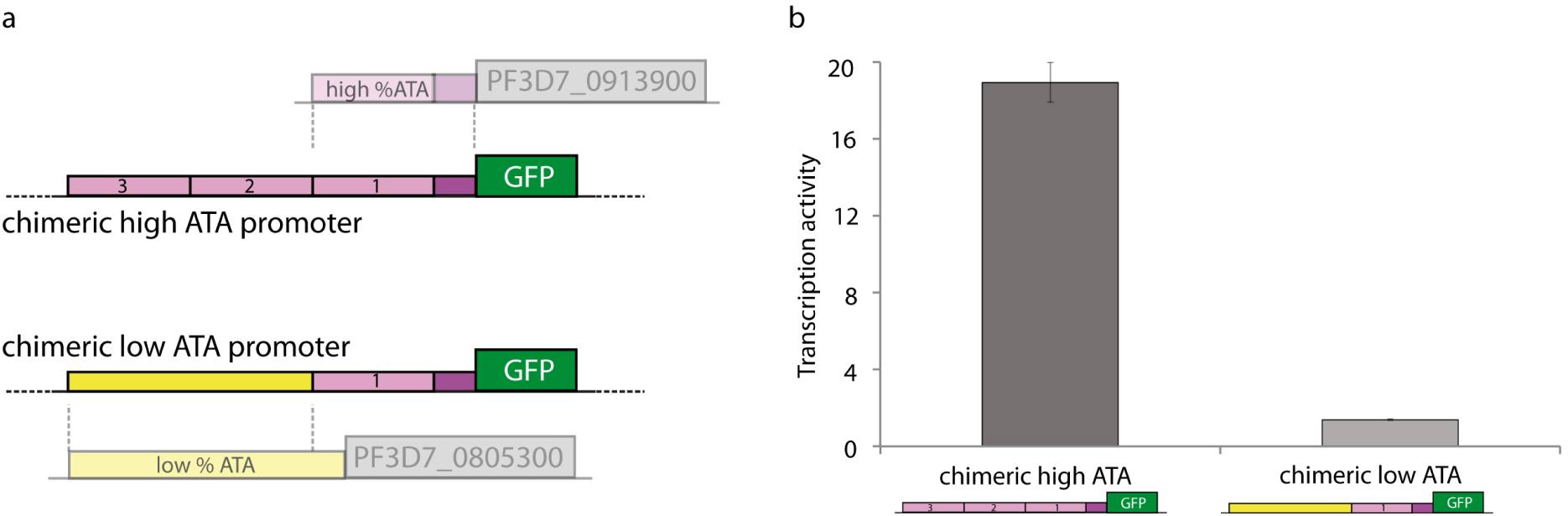
In vivo experimental validation in P. falciparum. **a** Schematic of the chimeric promoters used in our report assay to monitor promoter activity. **b** Transcriptional activity quatification by qPCR analysis of RNA collected at ring stages parasites. Here, one representative transgenic parasite clone. See Material and Methods for details.

## Discussion

Gene expression in Eukaryotes is orchestrated at different levels and by different mechanisms to ensure the wide variety of responses associated with the different cell types, stages and conditions. Besides traditional short TFBSs, long regions with specific nucleotide compositions may constitute another type of regulatory elements. While several *in silico* approaches exist for characterizing short TFBSs, to our knowledge, no methods dedicated to LREs have been proposed so far. We present in this paper a computational approach specifically designed to characterize long regulatory elements. Applied to various genomes and expression data, our method revealed that LREs seem to exist and to be active in a wide range of species and conditions. The nature of these regions greatly varies between species and, in some cases, between conditions, but, surprisingly, they seem to control a substantial part of gene expression in all studied organisms.

It is important to note that the apparent variety of LREs likely implies heterogeneous mechanisms of gene regulation. This is corroborated by our study of *P. falciparum* LREs, which showed that, depending on their location on the genome, they can be involved in transcriptomic or post-transcriptomic regulation mechanisms. Obviously, the exact nature of these mechanisms remains to be investigated and constitutes the main question raised by this study. LREs may constitute “loose” binding sites for certain DNA or RNA binding proteins as proposed by Quante and Birds (45), that may regulate for instance nucleosome occupancy (40), 3D genome architecture (42) and/or alternative splicing (46). DExTER provides a tool that will help design dedicated experiments aimed at better characterizing the contribution of LREs in these processes.

One striking observation is the distinct regulation dynamics in the different species. While regulation with LRE appears to be highly dynamic in *Plasmodium* species and *T. gondii*, in multicellular organisms these mechanisms seem to be much more static when different tissues are compared. One hypothesis could be that LREs in these species are used to fit a kind of rough, basal, transcription level for each gene. Adjustments to these basal levels could then be done in a tissue dependent way by other mechanisms that do not involve LREs. Interestingly, contrary to what is observed in tissues (with the exception of gonads), *Drosophila* and *C. elegans* embryo development also shows a dynamic behavior that goes along with a switch of the most important LRE positions. The gene body and the whole region seem more important at early time points, but they gradually loose their importance along the course of development in favor of central or 5’UTR regions.

In *P. falciparum*, LREs seem very important at every phase of the life cycle, and especially in IEC. Experiments with CNNs corroborated these results, although the accuracy achieved with these models was lower than with DExTER. Several reasons may explain the difference of accuracy, one reason being that the CNN architecture used here, which is classically used in regulatory genomics to identify TFBS motifs, may not be the best architecture to capture LRE features. Other architectures have indeed been proposed to predict transcription from CAGE data (28; 29). Hence, from a purely predictive perspective, it seems that some work is needed to propose an architecture capable of fully capturing LREs with CNNs. On the other hand, we must stress out that controlling what is exactly learned by CNNs is a difficult task (33). When used for modeling gene expression, these models likely capture a mixture of regulatory elements, including traditional TFBS motifs, LREs, and potentially many other kinds of regulatory elements. Because disentangling all these effects seems hazardous, we believe that direct approaches like DExTER constitute better alternatives than CNNs for studying the specific effect of LREs.

Our study also suggests that LREs are associated with transcriptomic and post-transcriptomic regulation in *P. falciparum*. LREs associated with nascent transcription are both upstream and non strand-specific, suggesting their involvement in transcriptomic regulation mechanisms, while LREs associated with mRNA stabilization are downstream and strand specific, pointing to post-transcriptomic regulation mechanism. There are several studies showing evidences for a control of gene expression at the post-transcriptional RNA level in this parasite. Lack of coordination between active transcription and mRNA abundance has been reported in *P. falciparum* (72; 73) and bioinformatics analysis have indicated that a significant percentage of Plasmodium genome encode RNA-binding proteins (4-10%) (74; 75).

Finally, in vivo analysis of the promoter activity of a chimeric DNA fragment showed higher transcription activity when the region was enriched in one of the LREs identified by DExTER as positively associated with RNA levels. It would be now interesting to investigate the molecular mechanism underlying the role of this LREs in gene regulation such as protein recruitment, nucleosome occupancy alteration or modulation of epigenetic marks.

Another question raised by this study is the reason for the apparent higher importance of LREs in *Plasmodium*, compared to other species. This might be linked to the paucity of TFs (2; 3) but also to the scarcity of distant regulatory sequences (enhancers) identified in *Plasmodium*, despite some work performed recently (76; 77). The potential regulatory mechanisms of short tandem repeats (78) may also explain the prominent role of LREs in *Plasmodium* gene regulation. For example, despite the existence of low proportion of TFs, the LREs may alter TFs binding, increasing their regulatory potential. Finally, LREs may also modulate expression changes through altering nucleosome positioning (79).

Another interesting observation is the prevalence of AT-rich k-mers in the identified LREs in *P. falciparum* genome. This may be somewhat surprising in a genome composed of more than 80% A+T, as any region is expected to be enriched for such k-mers. While this assertion is globally true, the frequency of a specific AT-rich k-mer in a specific region can nonetheless fluctuate between genes, and our study shows that these fluctuations are linked to gene expression. Moreover, we also showed that all AT-rich LREs have not the same correlation profile with gene expression (see Figure 7c). For example, ATAT [-1196,-126] has high correlation with expression in the IEC but lower correlation in gametocyte and ookinete stages, while TATAT [-1196,1925] shows moderate correlation in the IEC and higher correlation in gametocyte. Similarly, while the above mentioned AT-rich LREs are positively correlated with expression, ATTA [-1925,1925] appears to be negatively correlated with expression, especially in gametocytes and ookinetes. Hence, AT richness per se cannot be associated with a standardized global response but it seems on the contrary that *P. falciparum* has developed a subtle regulatory vocabulary largely based on these two nucleotides which, depending on the region and the exact k-mer, may generate different responses.

## Material and Methods

### The DExTER method

#### Step 1 - Feature extraction

We developed a procedure to identify pairs of (k-mer,region) for which the frequency of the k-mer in the region is correlated with gene expression. Starting from a 2-mer in the whole sequences, the procedure alternates two steps.

- The first step is the **segmentation** step (magenta arrows in the exploration graph of Figure 1). For this, sequences are first segmented in different bins defined from the alignment point (anchor). We used 13 bins in our experiments. The size of the bins are determined with the polynomial (*x*+*a*)^3^, with *x* being the rank of the bins with respect to the anchor (the bin centered on the anchor has rank 0, while bins immediately on the left and right hand sides of this bin have rank 1, etc.). *a* is a parameter determined automatically by the procedure in order to cover the whole sequences with the required number of bins (here 13) in the best possible way. With this method, bins close to the anchor are shorter than bins away from this point. When the binning is done, a lattice representing different regions that can be constructed from these bins is computed (see Figure 1). The top of the lattice represents the whole sequence, while lower nodes represent smaller regions. At each node, the correlation between the 2-mer frequency in the associated region and gene expression is computed, and the region with highest correlation is identified. If this correlation is sufficiently higher than the correlation associated with the top node (whole sequences) the region is selected. This procedure is resumed on all non-overlapping regions until the whole sequences are covered or the remaining correlations are lower than the correlation of the top node.
- Every identified region is then investigated for an **expansion** step of the 2-mer (green arrow in the exploration graph of Figure 1). Here, the goal is to identify (k+1)-mers whose correlation on the identified region is higher than the original k-mer. For this, the 8 possible (k+1)-mers obtained by concatenating a nucleotide on the left or right hand side of the k-mer are constructed. The correlations between gene expression and the frequency of these new k-mers in the region are computed, and k-mers that improve correlation are identified.

The whole procedure (segmentation + expansion) is resumed iteratively, until no improvement is observed. Then a new exploration starting from a different 2-mer is ran until every 2-mer has been explored. At each step of these explorations, regions and k-mers that improve correlations over the previous step (*i.e.* all nodes of the exploration graph of Figure 1) are stored and form the list of variables returned at the end of the procedure.

#### Step 2 - Feature selection and learning

Once all potential variables have been extracted, a regression model is learned and the best variables are identified. If only one gene expression data set is available, variable selection (Equation 1) is performed using the LASSO (Least Absolute Shrinkage and Selection Operator) (50): by penalizing the absolute size of the regression coefficients (l1-norm), the LASSO drives the coefficients of irrelevant variables to zero, thus performing automatic variable selection.

If several gene expression data are available for one species (as it is the case in the paper), we make use of multitask learning so that all models are learned simultaneously with a global penalization. Multitask learning exploits the relationships between the various learning tasks in order to improve inference performance. Here, in order to stabilize variable selection, we encourage that each feature is either selected in all samples or never selected. The group LASSO (80) is naturally suited for this situation. In particular, if the feature extraction step has identified the same k-mer in similar but slightly different regions in the different data, group LASSO encourages the selection of a common region for all models.

### Convolutional neural networks

We build a convolutional neural networks using the keras implementation (81) and with an architecture similar to the architecture proposed in DeepSea (25), *i.e.*:

- convolution layer (32 kernels, window size 8, step size 1, dropout 20%),
- max pooling layer (size 2),
- convolution layer (64 kernels, window size 8, step size 1, dropout 20%),
- max pooling layer (size 2),
- convolution layer (64 kernels, window size 8, step size 1, dropout 20%),
- max pooling layer (size 2),
- dense layer (64 kernels, dropout 50%),
- flatten layer,
- dense layer (1 kernel).

In each kernel we used the activation function relu and the glorot uniform initializer. The adam optimizer was used to minimize the mean squared error (MSE) loss function. 5% training sequences were used as validation sequences. We defined an early stopping condition on the MSE with a patience of 2. As for DExTER, 2/3 genes were used for training and the remaining 1/3 was used to measure the correlation between predicted and observed expression.

### Measure of variable importance

We devised an *ad hoc* procedure based on LASSO penalty and model error for measuring the importance of the different variables of a model. Given a penalization constraint *λ*, the LASSO procedure searches the model parameters that minimize the prediction error (MSE) subject to the constraint. In practice, a grid of constraints of decreasing values is initialized, and a model is learned for each value. The result is a series of models with increasing number of parameters. To identify the most important variables of a model in a given condition, we took the model with 15 parameters and estimated the importance of each of the 15 variables in the following way. Given a variable *X*, its importance was estimated by the MSE difference between the complete model and the model obtained by setting *β*_*X*_ to 0.

### Variable locations

Variables were assigned to 6 gene regions: distal and close promoters, central region, 5’UTR, gene body, and the whole region. For 5’UTR and gene body, the region boundaries were defined on the basis of the median size of the annotations found for each species. Here, the gene body region refers to the genomic region downstream of 5’UTR (it can potentially include introns). Note that for *P. falciparum* and *P. berghei*, the sequences are aligned on the AUG instead of the TSS, so the 5’UTR is defined upstream point 0. For regions different from 5’UTR and gene body (*i.e.* distal and close promoters, central and whole regions) boundaries were not based on existing genome annotations and the same values were used for all species. The table below reports the boundaries used for defining all gene regions in the different species:

| species | dist. prom. | close prom. | center | 5'UTR | gene body | whole |
| --- | --- | --- | --- | --- | --- | --- |
| *P. falciparum* | [-2000,-300] | [-300,-50] | [-500,500] | [-50,0] | [0,1715] | [-2000,2000] |
| *P. berghei* | [-2000,-300] | [-300,-50] | [-500,500] | [-50,0] | [0,1715] | [-2000,2000] |
| *T. gondii* | [-2000,-300] | [-300,0] | [-500,500] | [0,454] | [454,2000] | [-2000,2000] |
| *S. cerevisiae* | [-2000,-300] | [-300,0] | [-500,500] | [0,43] | [43,1014] | [-2000,2000] |
| Human | [-2000,-300] | [-300,0] | [-500,500] | [0,86] | [86,2000] | [-2000,2000] |
| *A. thaliana* | [-2000,-300] | [-300,0] | [-500,500] | [0,131] | [131,2000] | [-2000,2000] |
| *Drosophila* | [-2000,-300] | [-300,0] | [-500,500] | [0,94] | [94,2000] | [-2000,2000] |
| *C. elegans* | [-2000,-300] | [-300,0] | [-500,500] | [0,26] | [26,2000] | [-2000,2000] |

We used the Jaccard index for assigning each variable identified by DExTER to the gene region that most resembles it. Namely, given a variable region *R*1, we searched for the gene region *R*2 for which the ratio 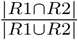 is the closest to 1.

### LRE conservation

The 10 most important variables of each species and conditions were identified and collected, and their correlations with expression were computed for every species and condition. For each variable, correlations were then normalized by conditions (z-score) to get the same range of values for each condition. Next, for each species and variable, we then identified and memorized the highest (in absolute value) normalized correlation found in any conditions of the species. We denote as 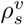 the best correlation found for variable *v* in species *s*. Then, at the level of the taxa, we used the minimum of the best correlations among all species of the taxa to measure the conservation of correlation of variable *v* (see Figure 6, down). For example, for the Apicomplexan taxa, we took the minimum of 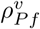, 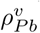and 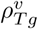 to assess the conservation of variable *v* at the level of Apicomplexan.

### Strand specificity

The strand specificity of a given variable for a given condition was measured on the basis of the correlation between the frequency of the variable in the different genes and the expression of the genes in the condition. More precisely, two correlations were computed and compared: the correlation computed on the frequencies measured on the plus strand (*ρ*_+_), and the correlation computed on the frequencies measured on the minus strand (*ρ*_−_). The quantity

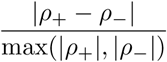

was then used to measure the strand specificity of the variable. With this measure, variables for which correlations are approximately the same on the two strands have strand specificity around 0, while variables with high correlation differences between strands have higher strand specificity. Note that this quantity is meaningless for the few k-mers for which the reverse complement is equal to the original k-mer (*i.e.* CpG, GpC, TpA and ApT for dinucleotides), because in this condition the k-mer occurrences are the same for both strands.

### Transcription *vs.* stabilization variables

In Painter et al. (2018) (15), the authors measured separately the level of nascent transcription and stabilized mRNA along the erythrocytic cycle. We ran DExTER on each available time point for these two types of data, and identified the 8 most important variables on each condition and time point, giving us a total of 15 different variables. For each variable, we computed its correlation with nascent transcription and with stabilized mRNA at each time point, and we summed the absolute value of theses correlations separately. This gives us two quantities for each variable: *ρ*_*v,trans*_ (resp. *ρ*_*v,stab*_) is the sum of the absolute value of correlations of variable *v* with transcription (resp. stabilization) at the different time points. Variables with *ρ*_*v,trans*_ > *ρ*_*v,stab*_ + 0.3 were associated with transcription, while variables with *ρ*_*v,stab*_ > *ρ*_*v,trans*_ + 0.3 were associated with stabilization. Variables with no clear differences were discarded.

### Data

- *P. falciparum*: RNA-seq data of the IEC were downloaded from the supplementary data of the original publication (51) and were log transformed. Life cycle RNA-seq data (68) were downloaded from PlasmoDB. Log transformed FPKM data were used for these analyses. Transcription *vs.* stabilization data (15) were obtained from the paper Supplementary data 1 and log transformed.
- *P. berghei*: RNA-seq data of the life cycle (52) were downloaded from PlasmoDB and log transformed.
- *T. gondii*: RNA-seq data of the life cycle (53) were downloaded from GEO (GSE108740), and the FPKM signal was log transformed.
- *S. cerevisiae*: RNA-seq data were were downloaded from GEO (GSE89554) and log transformed. Only the conditions (alpha, NaCl) in *S. cerevisiae* were used for the analysis.
- Human: For tissues, RNA-seq data were downloded from GTeX. We used the log of median TPM of 7 tissues for the analyses: Transformed fibroblasts, Esophagus - Mucosa, Lung, Minor Salivary Gland, Pancreas, Pituitary, Whole blood. For developmental series, we used data published in (60). Expression data were downloaded from GEO (GSE101571) and log transformed.
- *Drosophila*: For tissues, we used the data of (55). Expression data were downloaded from GEO (GSE99574, dmel.nrc.FB) and log transformed. Only the first biological repeat of each tissue was used in the analyses. For developmental series, we used the fly data produced in reference (56). Data were downloaded from GEO (GSE60471, DM) and log transformed. 10 time points along the whole time series were analyzed.
- *C. elegans*: For tissues, we used data from the cell atlas of worm (57). Data were downloaded from the Rdata file available on the cell atlas (http://atlas.gs.washington.edu/worm-rna/) and the log of the TPM of each available tissue were used for analyses. For developmental series, we used the data of published in reference (58). Data were downloaded from GEO (GSE87528, MA 20 strains) and log transformed. Only time points related to strain #550 were analyzed.
- *A. thaliana*: For tissues, we used the data published in ref. (61). Data were downloaded from ArrayExpress (E-GEOD-38612, FPKM) and log transformed. For development, we used the series published in (62). Data were downloaded from GEO (GSE74692, processed data) and log transformed. Only the first biological repeat of each time point of the WT series were used for the analysis.

### In vivo experimental validation in *P. falciparum*

#### Cloning of DNA constructs

All PCR amplifications were done with high-fidelity polymerase PfuUltra II Fusion HS DNA Polymerase (Agilent Technologies) following the recommended protocols, except we lowered the elongation temperature to 62°C. All cloning reactions used the In-Fusion ® HD Cloning Kit (Takara Bio USA, Inc.) and followed the manufacturer’s protocol. All PCR and digestions were purified using PCR Clean Up Kit (Macherey-Nagel) and followed the manufacturer’s protocol, except we performed 4-6 washes using 700*µ*L buffer NT3. All cloning and plasmid amplifications were done in Escherichia coli, XL10-Gold Ultracompetent Cells (Stratagene). All minipreps and maxipreps were performed using NucleoSpin Plasmid, Mini kit for plasmid DNA (Macherey-Nagel), and NucleoBond Xtra Maxi Plus kit for transfection-grade plasmid DNA (Macherey-Nagel), respectively. Sanger sequencing confirmed the absence of undesired mutations in the homology regions, the guide sequence, and the recombinant region.

The plasmids pBLD587_highATA_PfGFP and pBLD587_lowATA_PfGFP were constructed in multiple cloning steps from a pBLD587_HAtag-GFP backbone containing a Pfs47 homology region (Knuepfer et al., 2017). The 3’ UTR from *P. falciparum* HRPII was amplified from a PCC1 plasmid (82) and cloned using primers 1 and 2 and restriction sites SpeI and HindIII. The PfGFP was designed by codon optimization of the GFP using codon usage tables for *P. falciparum* 3D7 from Codon Usage Database http://www.kazusa.or.jp/codon, minimizing GU wobble pairings (83), and adding the 15-bp homology necessary for InFusion cloning. PfGFP was ordered as a DNA gBlocks® gene fragment from Integrated DNA Technologies and cloned using restriction sites HindIII and BamHI. Next, the core promoter and 356 bp DNA fragment that includes the identified ATA enriched region of PF3D7_0913900 was amplified from *P. falciparum* 3D7 genomic DNA and cloned using primers 3 and 4 and restriction sites XhoI and BamHI resulting the plasmid (pBLD587start). To generate the pBLD587_lowATA_PfGFP plasmid, we amplified 712 bp PF3D7_0805300 5’ intergenic region from *P. falciparum* 3D7 gDNA and cloned using primers 5 and 6 and restriction sites NotI and XhoI into plasmid pBLD587start. To generate the pBLD587_highATA_PfGFP plasmid, the second 356 bp region ATA enriched from PF3D7_0913900 was amplified and cloned it using primers 7 and 8 and restrictions sites AflII and NotI into pBLD587start plasmid. Next, the third 356 bp ATA enriched region was amplified from previous 3’ part of PF3D7_0913900 5’UTR amplicon and cloned using primers 9 and 10 and restrictions sites XhoI and AflII.

Primers:

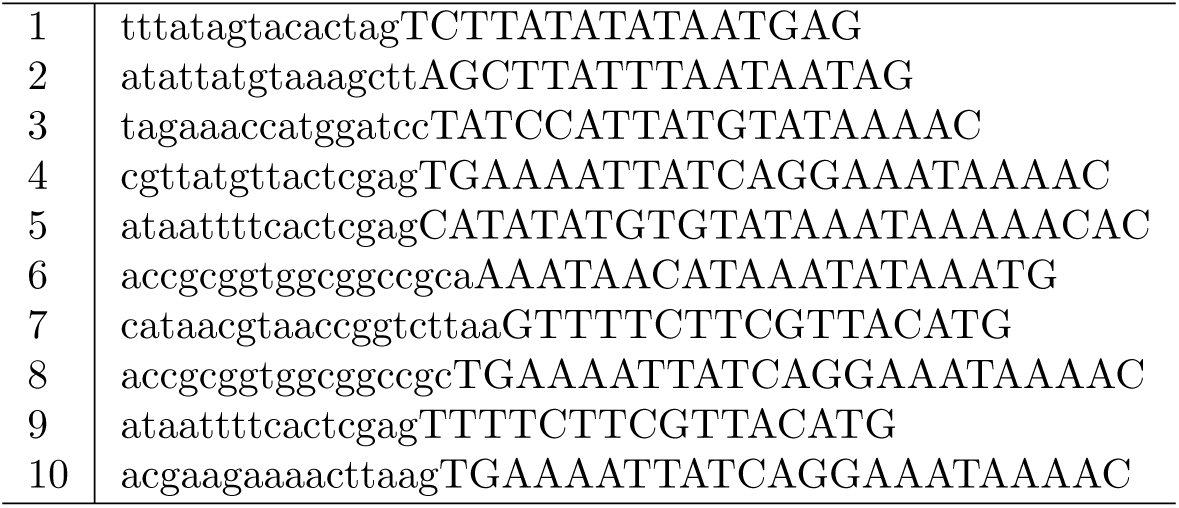

Nucleotides in lowercase show the overhangs introduced into oligonucleotides that are necessary to use InFusion cloning.

#### Parasite culture and transfection

*P. falciparum* 3D7 strain (MR4, ATCC) was cultivated in complete RPMI containing 5% human serum and 0.5% Albumax II (Thermo Fisher Scientific) and A-type human blood at 37°C 5% CO_2_, 5% O_2_ under agitation. Synchronous parasites were obtained by treating infected red blood cells with 9 volumes of 5% sorbitol for 10 min at 37°C. After one cycle, rings were used for transfections, using 60 *µ*g of each plasmid (pBLD587 highATA and pBLD587 lowATA) plus 60 *µ*g of pfs547 (kindly provided by Christiaan van Ooij), containing Cas9 and the Pfs47 guide RNA. Ring transfection was performed as published (84) with one pulse at 310 V, 950 *µ*F in a GenePulser Xcell (Bio-Rad) in 0.2 cm cuvettes (Bio-Rad). After electroporation, parasites were cultivated for one day without drug selection, followed by 5 days in which media containing 2.5 nM WR99210 was changed daily, and then every 2 days until drug-resistant parasites appeared in the cultures. Once integration was confirmed by PCR using primers checkPfs47F (CATTCCTAACACATTATGTGTATAACATTTTATGC) and checkPfs47R (CATATGCTAACATACATGTAAAAAATTACAATCAG), parasites were cultivated without drug pressure and cloned to obtain episome-free parasites.

#### qPCR analysis

For qPCR analysis, late-stage parasites were purified by gelatin flotation (Goodyer ID et al. 1994 Ann Trop Med Parasitol) and left to reinvade for 6 hours, after which a sorbitol treatment was applied to eliminate parasites that did not reinvade, allowing only young rings to continue the cycle. After one cycle, rings at 14 hours post-invasion were collected by 0.15% saponin lysis of RBCs, RNA was extracted by Trizol (Invitrogen), quantified by NanoDrop (Ozyme) and 1 *µ*g of total RNA was reverse-transcribed using Superscript IV (Invitrogen/Life technologies). Quantitative PCR was done in a LightCycler480 (Roche - Plateforme qPHD UM2 / Montpellier GenomiX) using Power SyBr Green PCR Master Mix (Thermo Scientific) and GFP primers (GFP_F1 TACCCAATG-TAATACCGGCGG, GFP_R1 GGTGACGGACCAGTTTTGTTG, GFP_F4 ACAAGAGTGTCTC-CCTCGAAC, GFP_R4 CCTGTGCCATGGCCTACTTTA), and as normalizers, seryl-tRNA synthetase (PF3D7_0717700), fructose-biphosphate aldolase (PF3D7_1444800) (85). Relative copy numbers were obtained from standard curves of genomic DNA of integrated parasites. Two clones were used for each transgenic parasite line.

### Availability of data and materials

The source code (python) of DExTER is available at address https://gite.lirmm.fr/menichelli/DExTER. This git repository also provides the R scripts for reproducing the main experiments described in the paper.

## Acknowledgements

We thank PlasmoDB for the invaluable malaria-database support, and Plateforme qPHD UM2 / Montpellier GenomiX for the Roche thermocyclers.

