## Supplementary Figures for "Identification of long regulatory elements in the genome of *Plasmodium falciparum* and other eukaryotes"

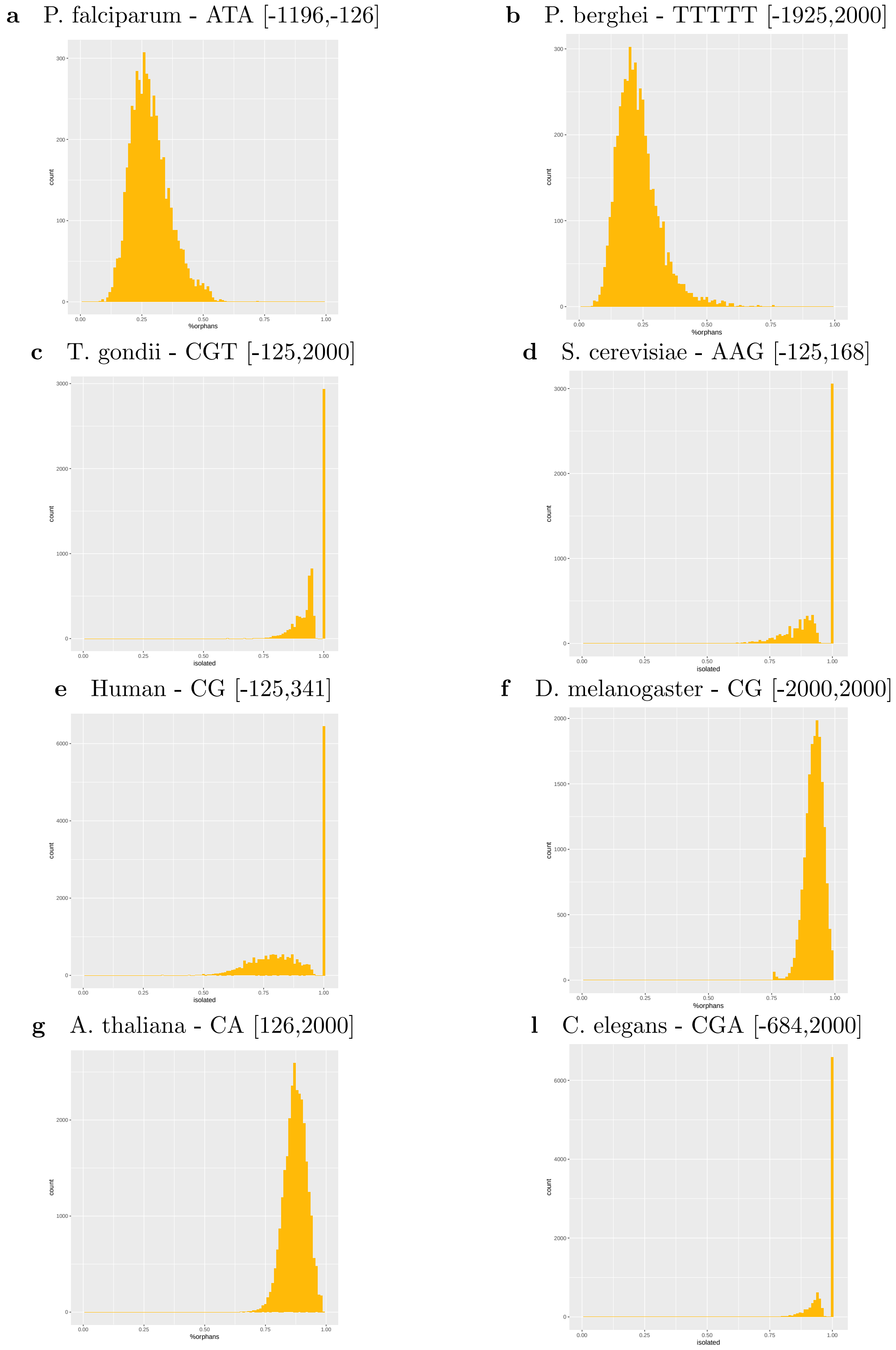


**Supp. Figure** 1: **Proportion of isolated occurrences of identified k-mers in regions.** These histograms report the proportion of isolated occurrences found in each gene of each species, for the most important variable of each species. A k-mer occurrence is considered as isolated if it is not immediately followed or preceded by another occurrence of the same k-mer. For these analyzes, k-mer repetitions can be either immediately consecutive or overlapping (for example, k-mer ATA is considered as being non-isolated in the two sequences ATAATA and ATATA).

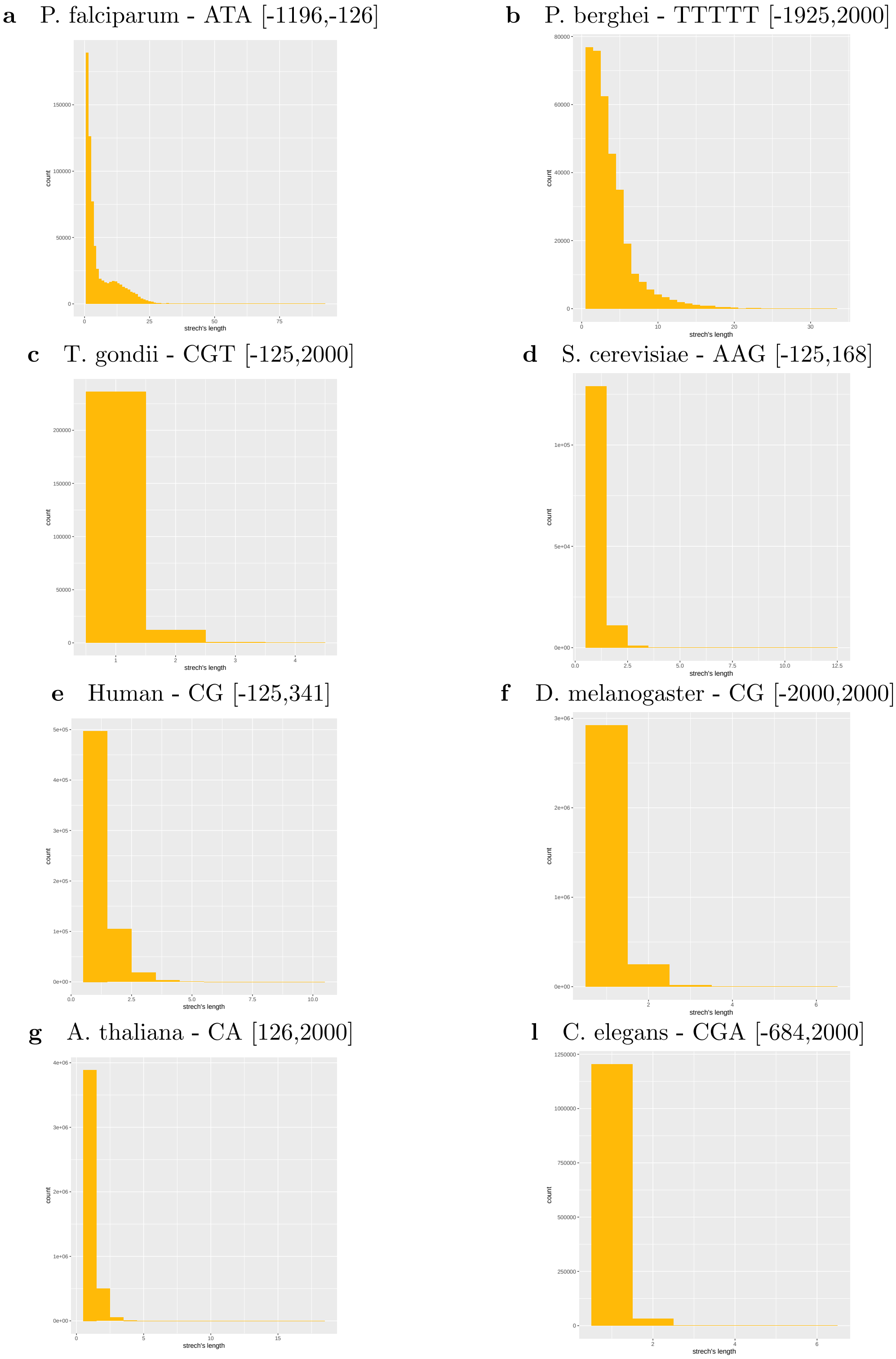


**Supp. Figure** 2: **Length of repetitive blocks.** These histograms report the length of the repetitive blocks in which the different k-mer occurrences are found. In this figure, isolated occurrences appears in "blocks" of length 1. As in Supp. Figure 1, k-mer repetitions can be either immediately consecutive or overlapping (for example, ATAATA and ATATA are tow repetitive blocks made up of two ATA occurrences each). Note that this figure report the length of the blocks in which each k-mer occurrence belong. This means that the same block appear as many times as the number of occurrences it contains.

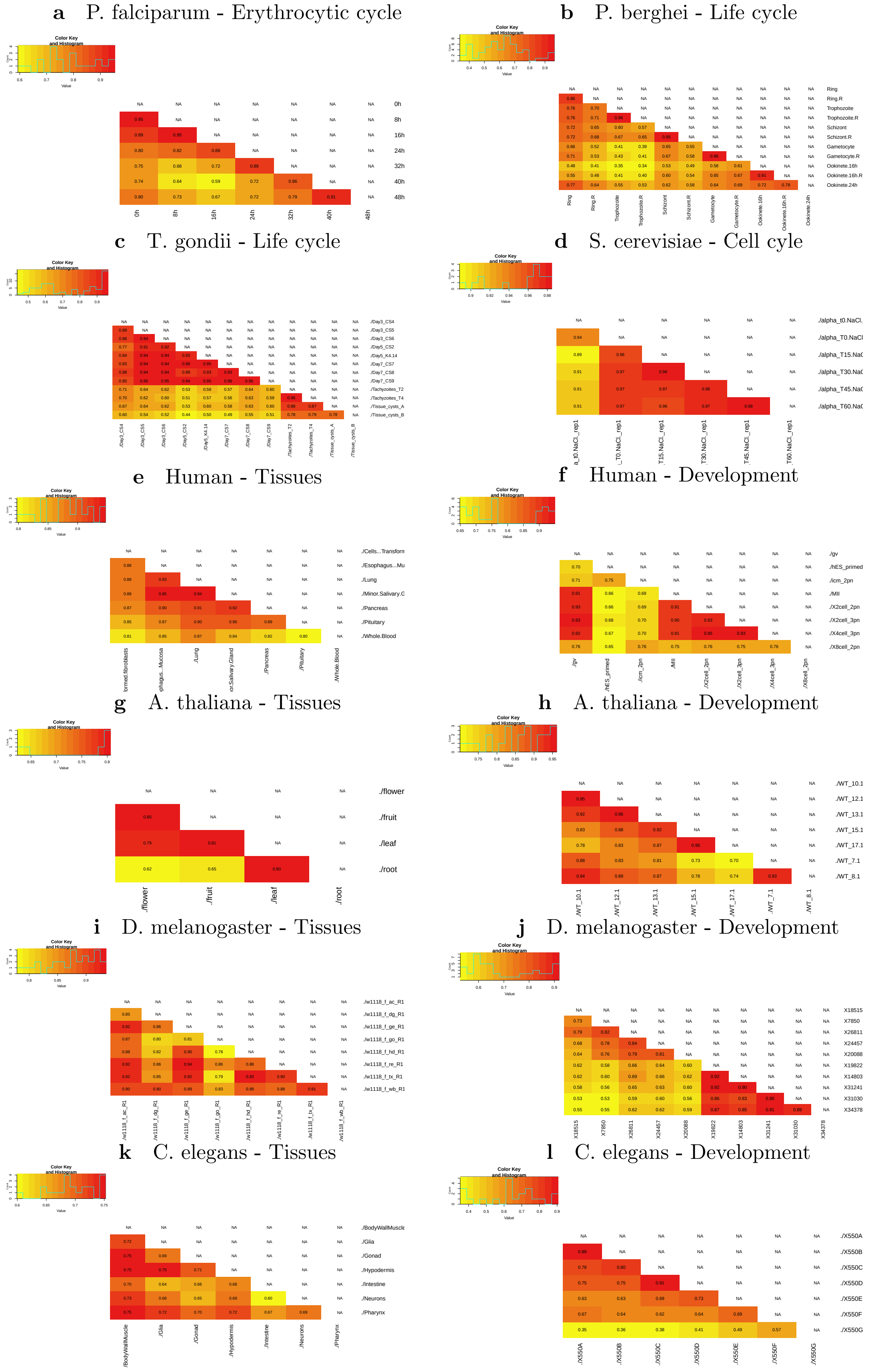


**Supp. Figure 3: Correlations between expression profiles of conditions.**

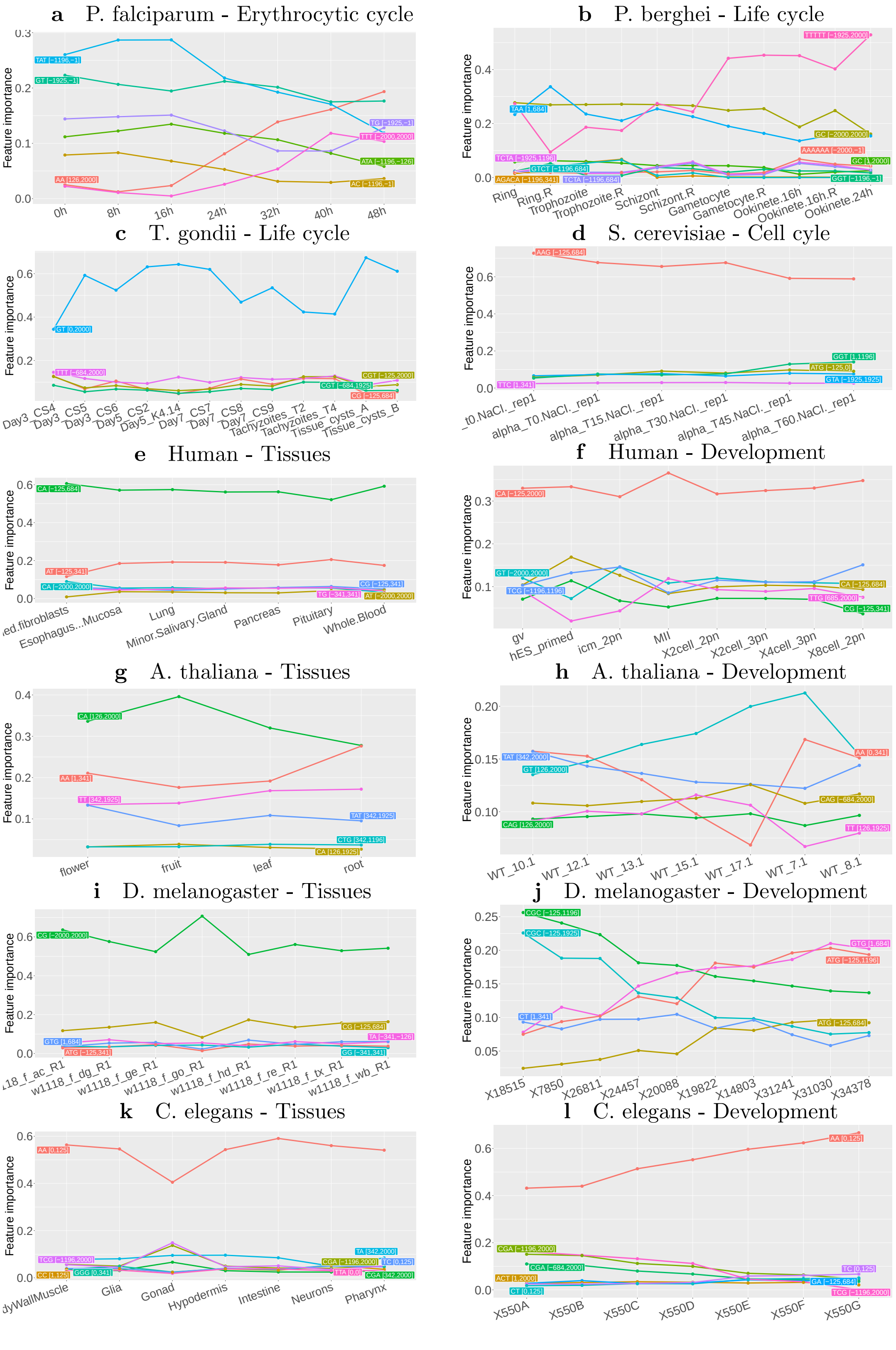


**Supp. Figure 4: Variable importance in the different models.** For each expression series, the 10 most important variables of each condition were identified, and their importance values were computed for all conditions of the series.

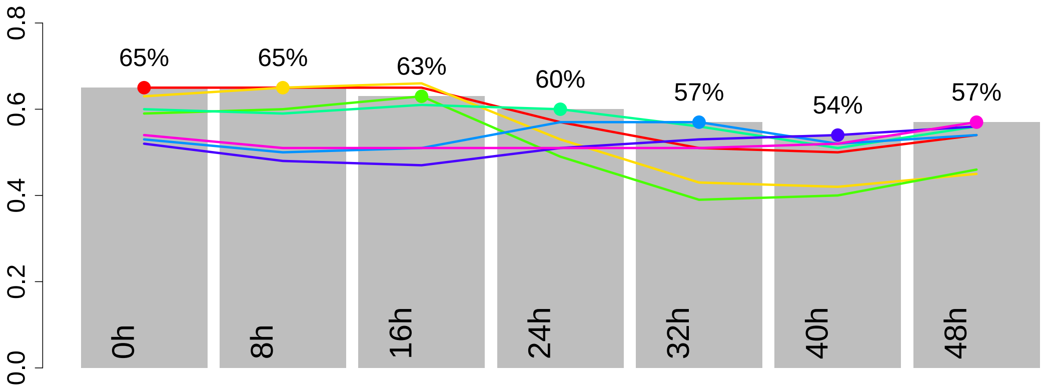


**Supp. Figure** 5: **Prediction of *P. falciparum* gene expression during erythrocytic cycle, with convolutional neural networks.** Grey charts represent the accuracy, measured as the correlation between predicted and observed gene expression, of the CNNs learned on different time points. Colored curves summarize the accuracy of a CNN learned on a specific time point (identified by a big dot of the same color) when used to predict the other time points of the erythrocytic cycle.

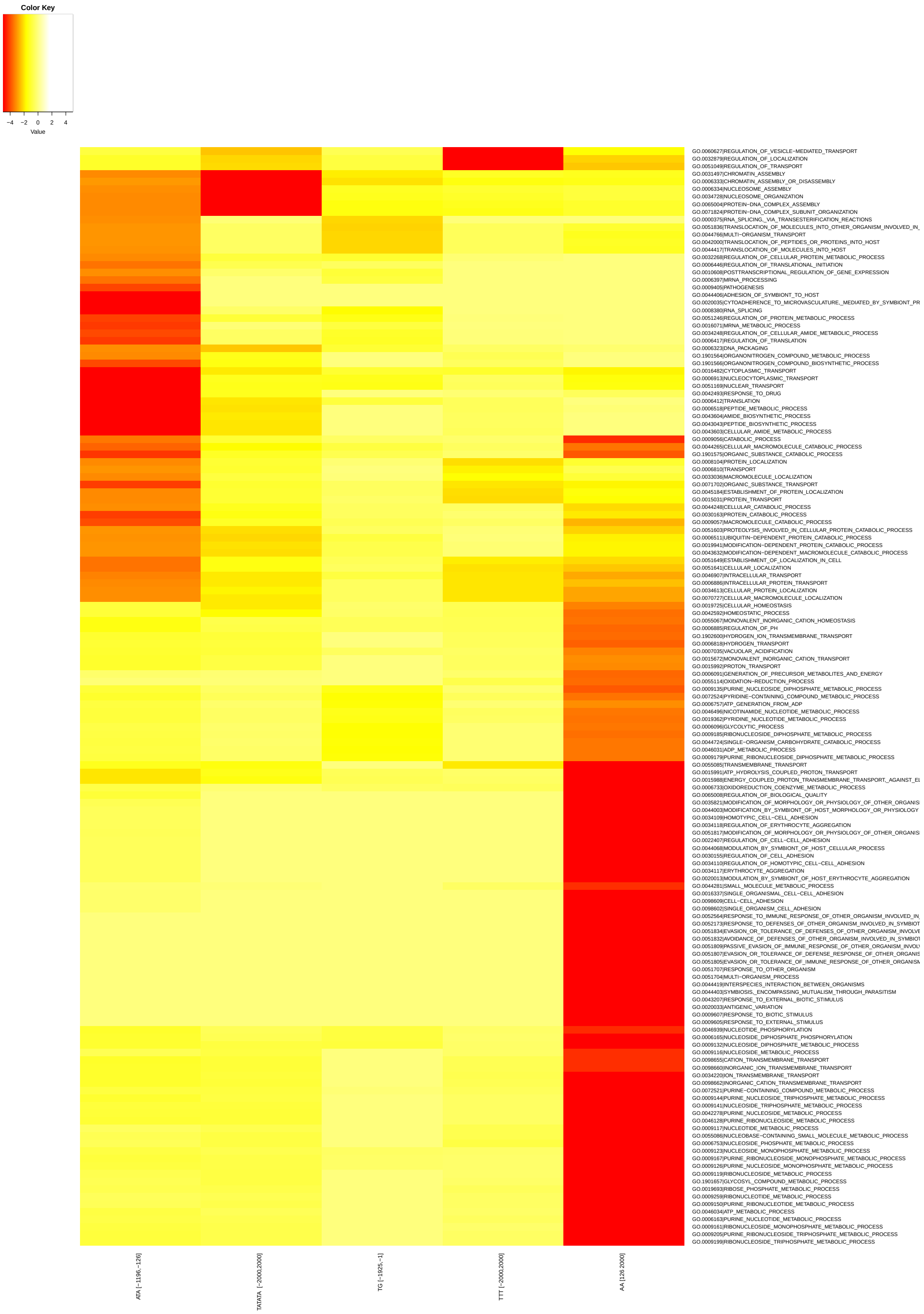


**Supp. Figure** 6: **Gene Ontology enrichment analysis of 5 important variables of *P. falciparum*.** The GSEA method [**?**] was used to identify GO terms enriched in genes associated with high ATA in region [-1196,-1], high TATATA in region [-2000,2000], low TG in region [-1925,-1], high TTT in region [-2000,2000], or low AA in region [126,2000]. The figure reports all GO terms associated with an FDR (q-value) below $10^{-3}$.

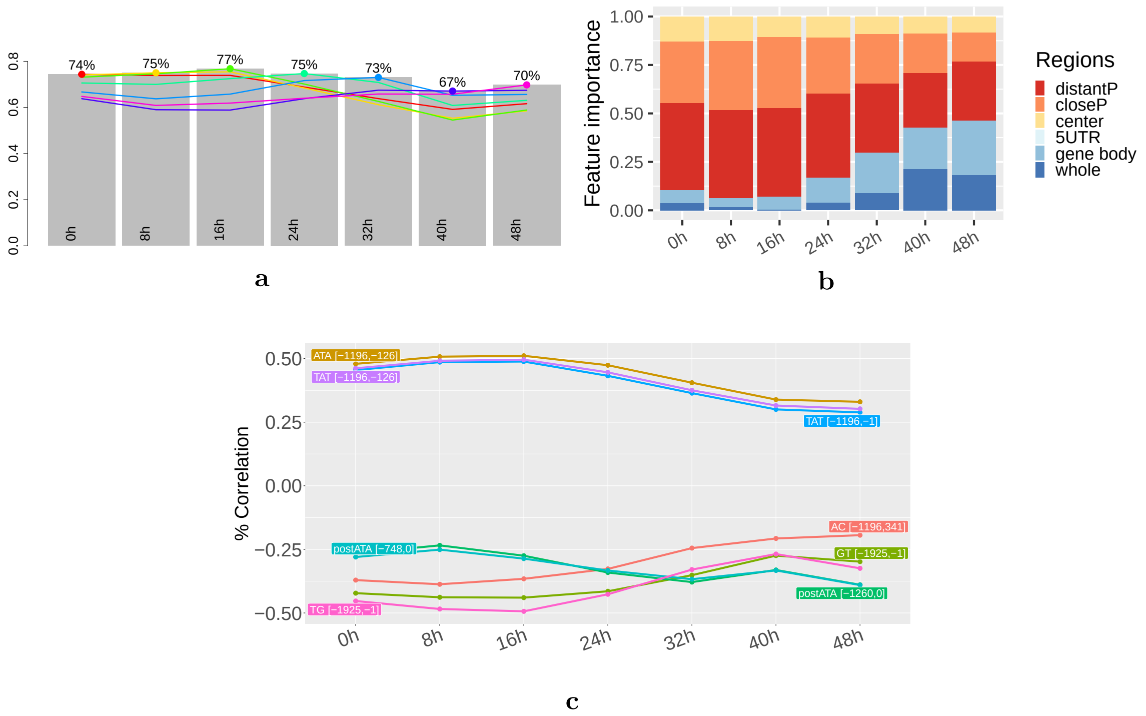


**Supp. Figure 7: Models incorporating cDNA variables - *P. falciparum* erythrocytic cycle.** These figures report the results achieved with models learned on all variables identified on genomic sequences (anchored on AUG codon) and on cDNA sequences (anchored on stop codon). **a** Grey charts represent the accuracy, measured as the correlation between predicted and observed gene expression, of the models learned on different phases of *P. falciparum* erythrocytic cycle. Colored curves summarize the accuracy of a model learned on a specific phase when used to predict gene expression of other phases. **b** Estimate of the importance of upstream, downstream, center, 5'UTR, gene body, and whole regions for predicting gene expression in the different phases. Variables identified on cDNA are classified in the gene body category. **c** Correlations between expression and k-mer frequency of the 10 most important variables identified at each phase. Variables identified on cDNA appears with prefix "post". On these variables, position 0 corresponds to the stop codon, while for the other variables position 0 corresponds to AUG codon.

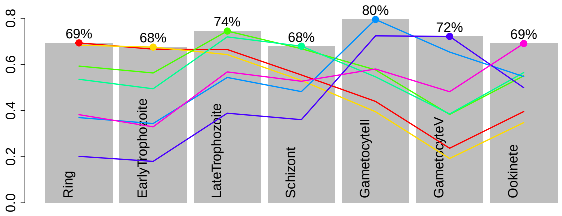


**a**

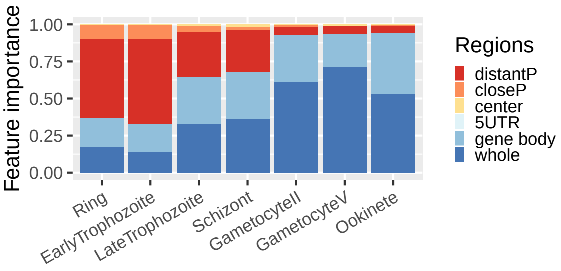


**b**

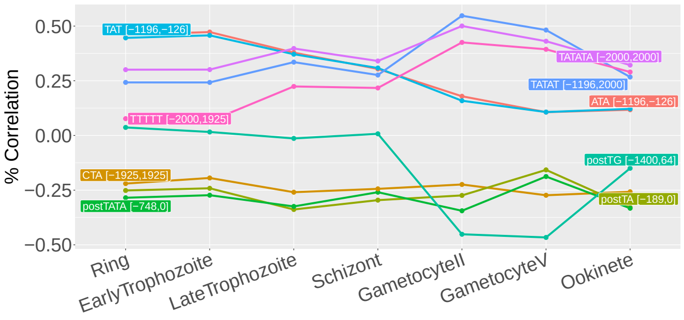


**c**

**Supp. Figure** 8: **Models incorporating cDNA variables - *P. falciparum* life cycle.** These figures report the results achieved with models learned when gathering all variables identified on genomic sequences (anchored on AUG codon) and on cDNA sequences (anchored on stop codon). **a** Grey charts represent the accuracy, measured as the correlation between predicted and observed gene expression, of the models learned on different phases of *P. falciparum* life cycle. Colored curves summarize the accuracy of a model learned on a specific phase when used to predict gene expression of other phases. **b** Estimate of the importance of upstream, downstream, center, 5'UTR, gene body and whole regions for predicting gene expression in the different phases. Variables identified on cDNA are classified in the gene body category. **c** Correlations between expression and k-mer frequency of the 10 most important variables identified at each phase. Variables identified on cDNA appears with prefix “post”. On these variables, position 0 corresponds to the stop codon, while for the other variables position 0 corresponds to AUG codon.

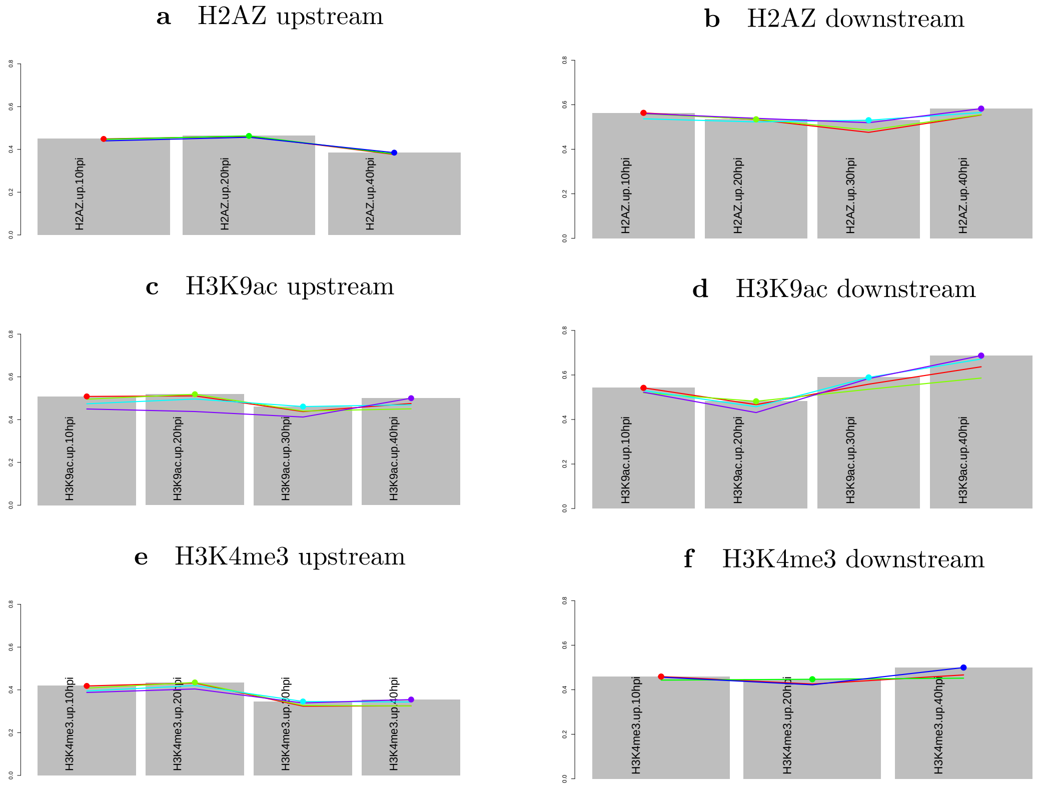


**Supp. Figure** 9: **DExTER accuracy for predicting H2AZ, H3K9ac and H3K4me3 histone marks.** Grey charts represent the accuracy, measured as the correlation between predicted and observed histone mark signal, on 4 time-points. Colored curves summarize the accuracy of a model learned on a specific time point when used to predict other time points of the same series.

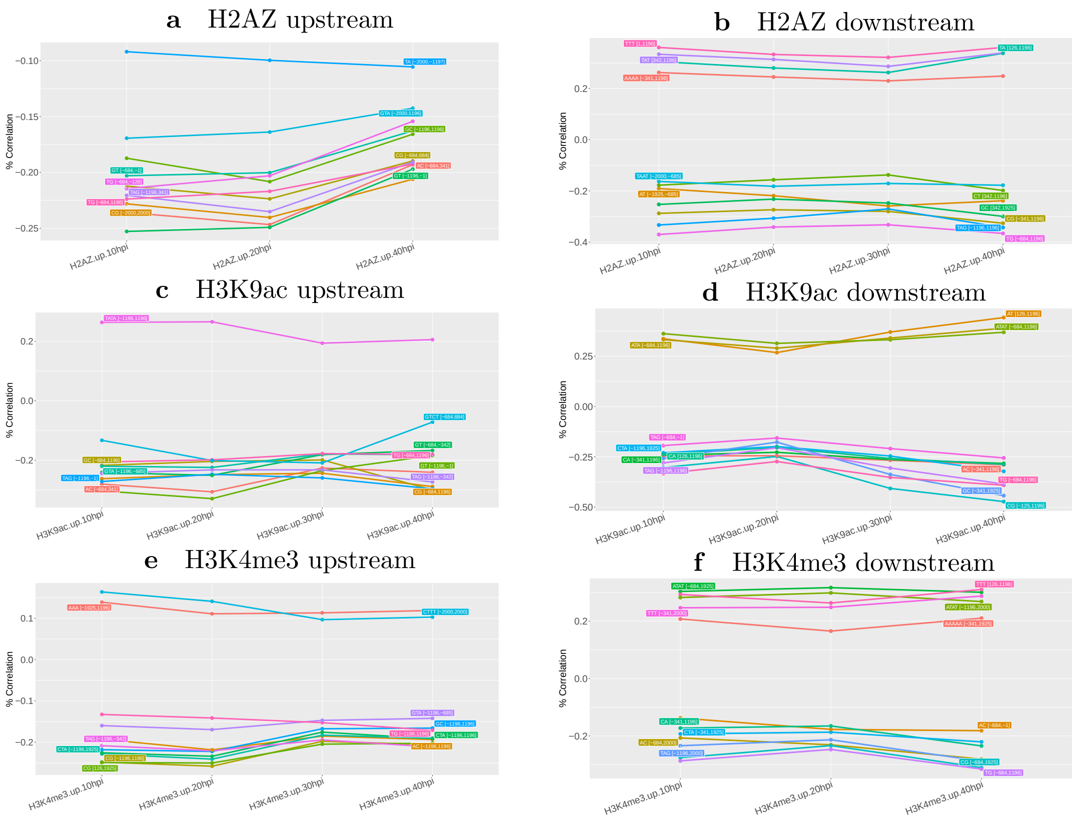


**Supp. Figure** 10: **Correlations between histone mark signal and k-mer frequency of the most important variables identified for the different histone marks and time points.** For each histone mark signal (upstream or downstream AUG), the 10 most important variables of each time points were identified, and their correlations to the histone mark were computed for all time points of the series.
